# Comparative Mucomic Analysis of Three Functionally Distinct *Cornu aspersum* Secretions

**DOI:** 10.1101/2022.11.16.516827

**Authors:** Antonio R. Cerullo, Maxwell B. McDermott, Lauren E. Pepi, Zhi-Lun Liu, Diariou Barry, Sheng Zhang, Xi Chen, Parastoo Azadi, Mande Holford, Adam B. Braunschweig

**Affiliations:** The Advanced Science Research Center, Graduate Center of the City University of New York, 85 St. Nicholas Terrace, New York, New York 10031, USA; The PhD Program in Biochemistry, Graduate Center of the City University of New York, 365 Fifth Avenue, New York, New York 10016, USA; Department of Chemistry and Biochemistry, Hunter College, 695 Park Avenue, New York, New York 10065, USA; Complex Carbohydrate Research Center, University of Georgia, 315 Riverbend Road, Athens, Georgia 30602, USA; Department of Chemical Engineering, The City College of New York, New York, New York 10031, USA; The PhD Program in Chemistry, Graduate Center of the City University of New York, 365 Fifth Avenue, New York, New York 10016, USA; The PhD Program in Physics, Graduate Center of the City University of New York, 365 Fifth Avenue, New York, New York 10016, USA; The PhD Program in Biology, Graduate Center of the City University of New York, 365 Fifth Avenue, New York, New York 10016, USA; Department of Invertebrate Zoology, The American Museum of Natural History, New York, New York 10024, USA

## Abstract

Every animal secretes mucus, placing them among the most diverse biological materials. Mucus hydrogels are complex mixtures of water, ions, carbohydrates, and proteins. Uncertainty surrounding their composition and how interactions between components contribute to mucus function complicates efforts to exploit their properties. There is substantial interest in commercializing mucus from the garden snail, *Cornu aspersum*, for skincare, drug delivery, tissue engineering, and composite materials. *C. asperum* secretes three mucus — one shielding the animal from environmental threats, one adhesive mucus from the pedal surface of the foot, and another pedal mucus that is lubricating. It remains a mystery how compositional differences account for their substantially different properties. Here, we characterize mucus proteins, glycosylation, ion content, and mechanical properties to understand structure-function relationships through an integrative “mucomics” approach. We identify new macromolecular components of these hydrogels, including a novel protein class termed Conserved Anterior Mollusk Proteins (CAMPs). Revealing differences between *C. aspersum* mucus shows how considering structure at all levels can inform the design of mucus-inspired materials.

## Main

Mollusca utilize mucus as glues,^1,2^ to create slick non-stick surfaces,^3,4^ and to facilitate innate immunity.^5,6^ The metabolic costs of mucus production can exceed one-quarter of mollusks’ energy budgets, indicating how important these materials are for survival.^7^ The structural component differentiating mammalian mucus from other soft materials are mucins — proteins containing densely *O-*glycosylated repetitive regions that form crosslinked networks from disulfide bonds, ion-bridges, and carbohydrate binding.^8,9^ However, molluscan mucus composition, and how they contribute to function, are not as well understood. Studies on *C. aspersum* mucus have focused on quantification of the protein within the mucus,^10^ bioactivity,^11^ the presence of antimicrobial peptides,^12^ or its ecological role.^13^ While mucins have been identified in aquatic snails and other mollusks,^14,15^ that contain the canonical A–B–A structure generally associated with mucins, with cysteine-rich (A) domains at the head and tail for disulfide bridging, and serine(Ser)/threonine(Thr)-rich (B) domains in the center possessing abundant glycosylation, no such mucins have been identified in *C. aspersum*.^8^ Rather, characterization of snail mucins has been limited to compositional analysis of amino acid and glycan residues, or studies on the molecular masses and hydrodynamic radii of the hydrogel particles.^16^ These studies found that proteins in the mucus secretions of *C. aspersum* contain overabundances of Ser/Thr residues, *N*-acetylgalactosamine (GalNAc), galactose (Gal), and fucose (Fuc) glycans, and the proteins had average molecular masses of 30 kDa, while mammalian mucins are typically 100 kDa to 1 MDa with an abundance of sialic acids.^17^ Broad proteomic analyses and profiling of snail mucus, focused on *C. aspersum* snail-snail signaling,^18^ microbial interactions,^19^ and comparison of proteins between multiple snail species.^20^ These studies illustrate that molluscan mucus contain proteins and glycans that are not found in mammalian mucus. Researchers have also investigated the role of ions in snail mucus and correlated increased CaCO_3_ content with increased mucus aggregation and adhesion.^3,21,22^ Notably, these prior studies analyze crude mucus collections, rather than purified mucus samples that reflect the protein compositions of the gels, themselves.

Despite these efforts, it remains unclear how differences in protein structure, ion concentration, glycosylation, and other factors operate synergistically to account for the substantial diversity in mucus material properties.^23^ Here, we apply a systematic comparative mucomic analysis — defined as the combination of genetic, chemical, and material studies to understand the structure-function relationships of mucus — of adhesive,^22^ lubricating,^3^ and protective^24^ mucus isolated from *C. aspersum*, which are named in accordance with the materials’ ecological function (Figure 1a). Transcriptomic and proteomic sequencing identified the proteins expressed in each mucus and their abundances. Glycomic mass spectrometry was then employed to identify the structures of the glycans decorating these proteins. Elemental analysis through scanning electron microscopy (SEM) coupled with energy dispersive X-ray spectroscopy (EDX) measured concentrations of various ions in the materials. Atomic force spectroscopy quantified the mechanical properties (elastic modulus, *E*, and work of adhesion, *W*) of the three samples. Comparison of these datasets reveal how *C. aspersum* exploit differential protein expression — including a series of previously uncharacterized proteins — glycosylation, and ion concentration are used to explain how these hydrogels behave as adhesives, lubricants, or protective barriers^25^ (Figure 1b).

**Figure 1.**
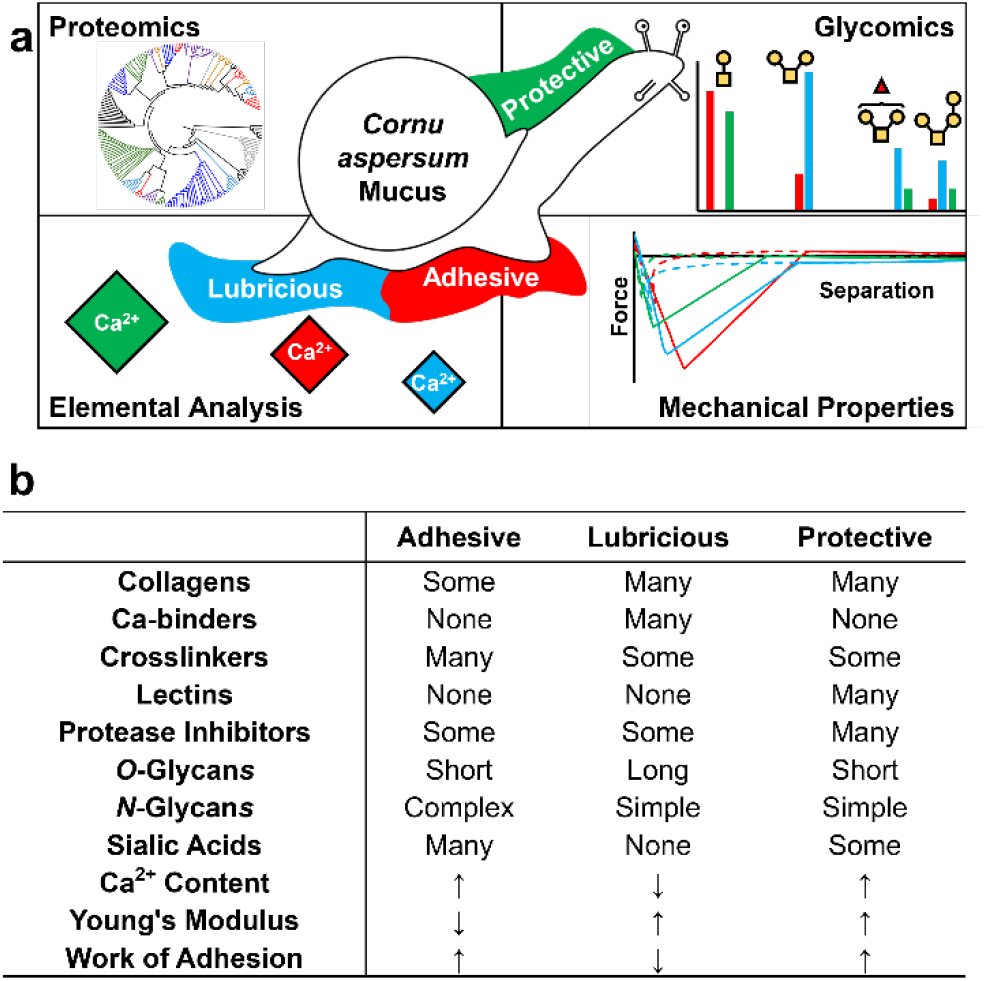
Mucomic analysis of *Cornu aspersum* snail mucus composition. a) Adhesive, lubricating, and protective mucus are subjected to an omics-style analysis to understand the composition and properties. b) Comparative overview of the compositions and properties of *C. aspersum* adhesive, lubricious, and protective mucus.

## Results

### Collection and purification of adhesive, lubricious, and protective snail mucus secretions

Adhesive, lubricious, and protective mucus samples were separately collected from *C. aspersum* snails (Supplementary Figure 1).^26^ The snails were placed onto inverted petri dishes, to which they attached, resulting in the deposition of adhesive mucus from the pedal surface of the foot. Lubricating mucus was collected from the trails left behind by the pedal surfaces of snails that had crawled along petri dishes. Protective mucus was scraped from the dorsal surface of the snail. Proteins embedded within the mucus gel were fractionated from cellular detritus so that analysis focused upon the components integrated into the mucus (Supplementary Figure 2). Isolated mucus all occurred as flocculent, beige substances with the consistency of cotton candy (Supplementary Figure 3). Purified mucus proteins were resuspended and subjected to spectrophotometric analysis to quantify the yield of protein samples at every purification step (Supplementary Figure 4, Supplementary Table 1). Initial protein concentrations in the solution of resuspended crude mucus were approximately 77 – 170 mg/mL for all three samples. Following purification, 3.9 – 7.4 % of the initial total protein was recovered.

### Identification and sequence alignment of snail mucus proteins

Shotgun proteomic sequencing supported by a *de novo* assembled transcriptome identified proteins in the purified mucus samples.^27^ As *C. aspersum*’s genome has not yet been sequenced, a transcriptomic reference database of actively translated genes found in mucus-producing tissue was produced from RNA extracted from the foot and back tissue of whole *C. aspersum* snails. From the 179,552 transcripts, 71 provided coding sequences for proteins based on the standard criteria of having a minimum of two identified peptides and a false discovery rate of less than 1.0 %.^18,28^ All proteins were quantified based on MS/MS proteomic abundance (Supplementary Figures 5–7, Supplementary Table 2). Many of the proteins had sequence similarity (*E*-value < 10^‒ 4^) to known proteins from snails or other mollusks, and were thus assigned identities corresponding to the specific proteins with which they shared highest similarity (Supplementary Table 3). More than 20 % of the proteins in each mucus matched proteins from other snails that have no reported function, referred to herein as “Snail” proteins, or appeared to be completely unique, which are referred to as “Novel” proteins. To better understand the relationships between our sequenced proteins and those of other mollusks, and to determine the functions of the identified and unidentified proteins, global alignment analysis was employed to cluster the genes by sequence similarity (Figure 2, Supplementary Figure 8). These clusters were assigned broad categorizations as inhibitors, enzymes, mucins, matrix, network, lectins, or ion-binders. Notably, there were two mucin clades — one large cluster of mucins broken into three smaller clades, and one separate smaller clade. Using this approach, the 12 Snail proteins and the 6 Novel proteins were each grouped into one of the aforementioned categories, allowing putative identifications that could not be made by BLASTP searches (Supplementary Table 4).

**Figure 2.**
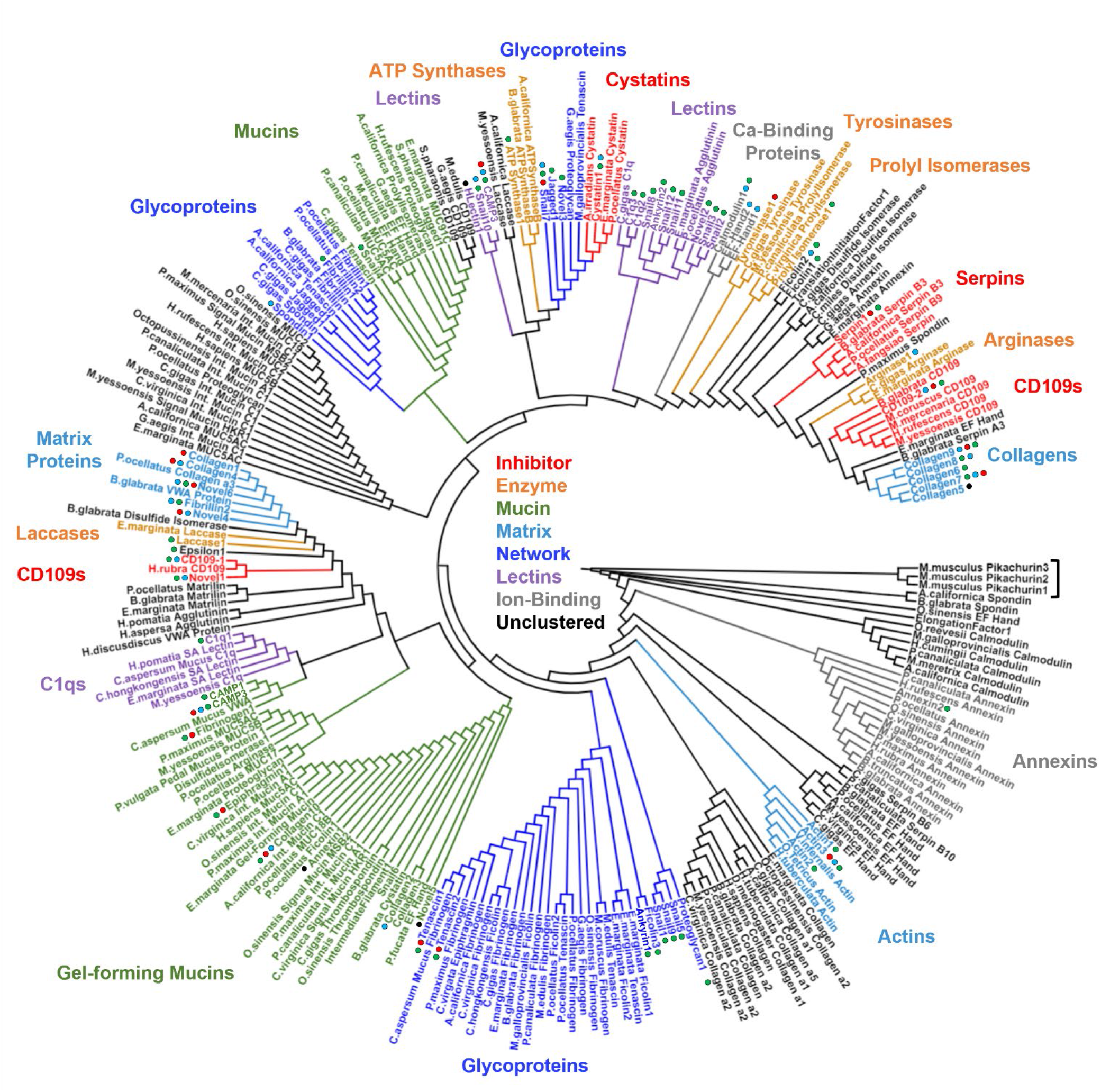
Dendrogram of snail mucus proteins based on sequence similarity. Clusters are colored according to protein function. The “Unclustered” (black) classification indicates a clade that had no discernible function or only contained reference proteins. Proteins identified in this study are labelled with circles. Circle color indicates the protein was found in adhesive (red), lubricating (blue), or protective (green) mucus. An outgroup, three *Mus musculus* proteins (Pikachurin1, Pikachurin2, Pikachurin3), is marked with a bracket. Dendrogram with branch lengths and included species is shown in Supplementary Figure 8.

### Characterization of adhesive, lubricating, and protective *C. aspersum* mucus proteins

Several glycoproteins that are components of oligomeric networks (independent of conventional extracellular matrix proteins), such as fibrinogens, ficolins, and tenascins, were identified, suggesting a wide diversity occurs in the protein-protein networks. These proteins could, in turn, cause differences in the mechanical behavior of these mucus.^29^ These glycoproteins make up about 15 % of the adhesive and protective mucus, and 5 % of the lubricating mucus, respectively. Importantly, BLASTP searches of several proteins returned A–B–A mucins as positive hits, which had not been identified in previous studies. It should be noted that mucins are challenging to identify via shotgun proteomics because their dense glycosylation limits enzymatic digestion,^30^ or via transcriptomics because of their tandem repeats^31^. A jagged-1-like protein (Jagged1), which is involved in extracellular signaling pathways,^32^ had sequence similarity to MUC2 from *Pygocentrus nattereri* (Red-bellied piranha). A spondin-like protein (Spondin1), which mediates cell-extracellular matrix interactions,^33^ also displayed similarity to MUC2, MUC5AC, MUC12, MUC16, and MUC19 from mollusks and other marine life. Spondin1’s sequence features several short regions that are either Ser- or Thr-rich. Curiously, these regions alternate between being Ser-rich and Thr-rich, meaning each region only incorporates one of these two amino acids. This protein is also 12% Cys by composition, which is more than five times greater than the natural abundance of Cys in invertebrate proteins,^34^ suggesting it has a propensity to form disulfide bridges. It is likely Spondin1 is a *C. aspersum* mucin because it contains repeating Ser/Thr-rich regions for potential *O*-glycosylation sites, similar to ‘B’ domains of mucins, and Cys-rich regions which can multimerize the protein by functioning like mucin ‘A’ domains. Vertebrate SCO-spondins, which are repetitive, highly glycosylated, and bind Ca^2+^, and are known orthologs of invertebrate mucins.^35^ From the alignment analysis, several proteins clustered within the large mucin families, including ones whose functions were not determined initially by BLAST. Snail6, Snail10, and Novel5 were clustered with mucins. Additionally, Snail1, Snail5, Snail9, and Novel3 fell within glycoprotein groupings. Thus, it appears that these secretions involve a combination of mucin-like proteins. Epiphragmin was identified in the adhesive mucus, which is used to create the epiphragm — the persistent glue that maintains bonds between the snail’s shell and substrate.^36^ This protein has been found previously in other snail species, such as the vineyard snail, *Cernuella virgata*, and is localized to the pedal surface of the foot,^18^ suggesting it is conserved in mollusks and has important function in snail adhesion. Notably, 30% of the adhesive mucus is composed of only two tenascin glycoproteins.

Extracellular matrix proteins comprise 40 – 50 % of all three *C. aspersum* mucus protein samples, with lubricating mucus incorporating more matrix proteins (50 %) than the other two (40 % each). Eleven unique collagen genes were identified, which were found previously to be expressed in snail mucus.^37^ While many of the matrix protein genes code for collagens, there are stark differences in abundances of these collagens between the samples. Collagen2, Collagen3, and Collagen11 are exclusive to lubricating mucus, while Collagen4, Collagen9, and Collagen10 are in all three mucus. Collagen7 and Collagen8 are more abundant in protective mucus than the other two. Collagen6 is exclusive to protective mucus. Collagen1 is found exclusively in adhesive mucus, albeit with very low abundance. Adhesive mucus shares Collagen4, Collagen7, Collagen9, and Collagen10 with the other mucus, however they less abundant in adhesive mucus than the lubricating and protective.^29^

Several enzymes that were found are involved in protein crosslinking, mucus network formation, or constructing biological glues, and likely serve a similar role in *C. aspersum* mucus.^38,39^ Cysteines are abundant in mucins, and disulfide isomerases, like the identified DisulfideIsomerase1, construct mucus gels by catalyzing interchain disulfide bonds.^39^ A prolyl isomerase, ProlylIsomerase1, was found, which has signaling and immune functions in mucus,^40^ and this class of proteins also regulates collagen crosslinking.^41^ A tyrosinase, Tyrosinase1, was found exclusively in the adhesive mucus, which catalyzes the formation of L-DOPA from tyrosine. As L-DOPA is involved in forming strong glues in *Perna viridis* mussels,^42^ this observation suggests that *C. aspersum* land snails may use a similar adhesive mechanism as marine mollusks.^42^ Tyrosinases also produce melanins, which are crosslinked networks formed through polymerization of phenolic molecules, and enzymes involved in melanin biosynthesis have been reported previously in snail and other invertebrate mucus secretions.^43^ Additionally, a laccase, Laccase1, was identified, which catalyzes the oxidation and crosslinking of phenolic compounds.^38^ Thus, tyrosinases, laccases, and other phenoloxidases may increase snail mucus integrity by crosslinking phenols of proteins and metabolites, similar to mechanisms used in other mollusks.^44^ No proteins strongly identifying with *P. viridis* mussel foot glue proteins were identified in this study, though Fibrillin1 and Fibrillin2 showed some sequence similarity to mussel foot proteins.^42^

Several proteins were found that have potential roles in defense. All mucus include 1 – 5 % protease inhibitor proteins, which can have infection-mitigating effects and also protect the protein scaffold from degradation.^45^ Mucins and other mucus proteins are Ser- and Cys-rich, thus the serine (serpins and CD109s)^46,47^ and cysteine (cystatins)^48^ protease inhibitors identified in this analysis could prevent pathogens from degrading the mucus barrier. While serine protease inhibitors were found in all mucus, the adhesive mucus did not contain any cysteine protease inhibitors. Protective mucus is 10 % lectins, which confer immune function in mollusks by protecting the snail’s skin from pathogens.^5^ The other two mucus were ∼2 % lectin. C1q lectins, which have immune function, complex with antigens, and noncovalently crosslink mucus glycoproteins,^49^ and Gal-specific H-type lectins,^50^ were exclusively found in the protective mucus. The protein C1q1 made up 6 % of the protective secretion and < 2 % of the other two.

Calcium ion (Ca^2+^)-binding proteins were entirely absent from the adhesive mucus and were minimally present in the protective mucus, but were abundant (∼10 %) in the lubricating mucus. This class of proteins includes annexins, calmodulins, and EF-hand proteins.^51^ Since these proteins were mainly found in the lubricating mucus, they may play a different role outside of forming gel networks. These proteins are involved in Ca^2+^-dependent signaling pathways, suggesting they relay environmental information back to mucus-producing tissue.^52^ Prior research has demonstrated that Ca^2+^ crosslink mucus gel particles.^3^ Thus, the presence of Ca^2+^-binders suggests that Ca^2+^ may have a different fate in the lubricating mucus than in adhesive and protective. It is possible these proteins function as ion traps, preventing the Ca^2+^ from participating in ion bridges between highly glycosylated mucus proteins.

For each mucus, 20 – 40 % of the identified proteins fell into the Snail and Novel groupings, meaning they had no identifiable function from BLASTP searches. Several of these proteins individually made up an appreciable amount of the mucus secretions. Over 40 % of the adhesive mucus’ is made of only 3 proteins, Snail7, Novel5, and Novel6. Novel5 comprised 7 % of the adhesive mucus, where it was exclusively found. Snail7 and Novel6 were shared across all three mucus, but made up a much greater proportion of the adhesive sample than the other two. Using alignment analysis, all 18 of the Snail and Novel genes clustered into clades and assigned putative functions (Supplementary Table 3). Novel1 was grouped with CD109 proteins. Novel5 clustered with gel-forming mucins. Novel6 clustered within a family of mollusk glycoproteins and Von Willebrandt Factor A (VWA) proteins. Snail6, which was abundant primarily in lubricating mucus, was clustered into the same gel-forming mucin family. Snail4 was also placed with mucins. Snail7 fell within a clade of mollusk glycoproteins. Snail1, Snail5, Snail9, and Novel3 were clustered into mollusk glycoprotein clades. Snail2, Snail3, Snail8, Snail10, Snail11, Snail12, and Novel2 were grouped with lectins.

### CAMPs, a new class of mollusk proteins

Three proteins, CAMP1, CAMP2, and CAMP3, were identified that could not be readily categorized into the aforementioned groups. Upon multiple sequence alignment and BLAST and HMMER searches, these CAMPs were found to share sequence identity to each other and previously found, but not well-characterized, mollusk proteins.^53^ These mollusk proteins from the databases contain lectin, VWA, and fibrinogen domains. The database proteins also share a general architecture with the three CAMPs identified here (Figure 3a, Supplementary Table 5). We deem this class of proteins as ‘CAMPs,’ or <u>C</u>onserved <u>A</u>nterior <u>M</u>ollusk <u>P</u>roteins because their *N*-terminal regions were nearly identical, but had entirely different *C*-terminal regions (Figure 3b). These proteins’ *N-*termini were abundant in Ser/Thr for potential glyscosylation, had Ca^2+^-binding pockets, and had oligomerization domains^54^ that are irregularly spaced throughout the protein sequences. The *C*-terminal regions of CAMP1 and CAMP3 were fibrinogen-like domains and CAMP2 contained a Gal-specific lectin domain. From sequence alignment analysis, CAMP1 and CAMP3 were clustered with *C. aspersum* mucus VWA protein and a fibrinogen. CAMP2 was clustered separately, as it was paired alongside a possible H-type (GalNAc-specific) lectin. It is possible the *N*-terminal domains are involved in physically integrating the proteins into the mucus gel through noncovalent linkages, such as H-bonds, disulfide, and ion bridges with mucin proteins, while the *C*-terminal domains provide protein functionality.

**Figure 3.**
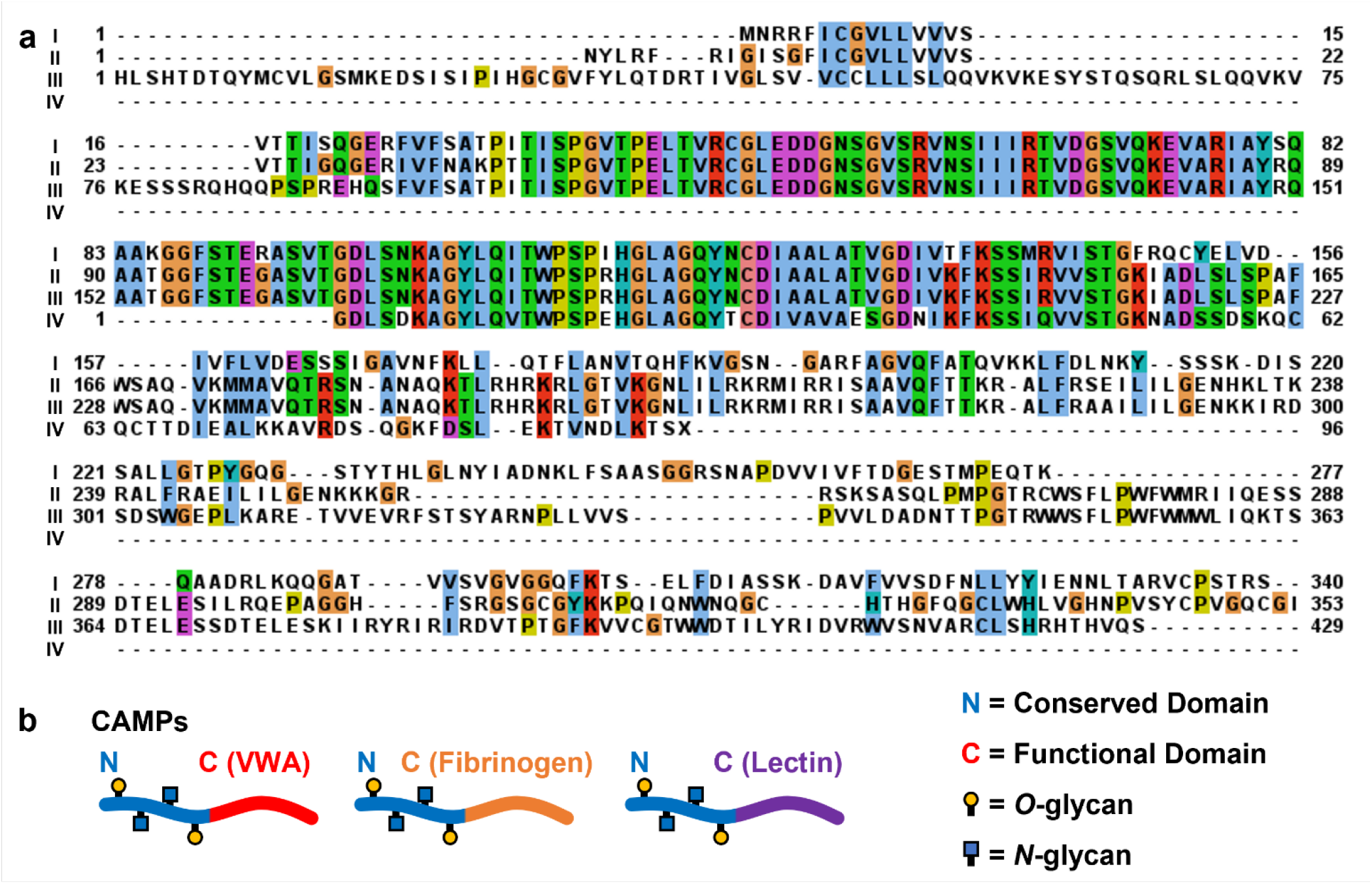
Sequence analysis of CAMPs reveals similarity among *N*-terminal domains with interchangeable *C*-terminal functional domains. a) Multiple sequence alignment between I) *C. aspserum* mucus VWA protein, II) CAMP1 (truncated at *C*-terminus), III) CAMP2, IV) CAMP3. b) Schematic of CAMP architectures, showing conserved *N*-termini but varied *C*-termini between proteins.

### Glycomic analysis of *C. aspersum* mucus

*O*-glycans in mucins, which are *O*-linked to Ser or Thr via a GalNAc residues, have been associated with lubrication, biological recognition, and network formation.^55^ As such, the *O*-glycan compositions of the three mucus were individually analyzed. *O*-glycans from the snail mucus proteins were extracted by *β*-elimination with sodium borohydride,^56^ and their structures and abundances were identified using permethylation and MALDI (matrix-assisted laser desorption/ionization) mass spectrometry (Supplementary Figures 9–11, Supplementary Table 6, Figure 4a).^57^ Experiments were conducted using both iodomethane and iodomethane-D3 to identify native methylation.^58^

**Figure 4.**
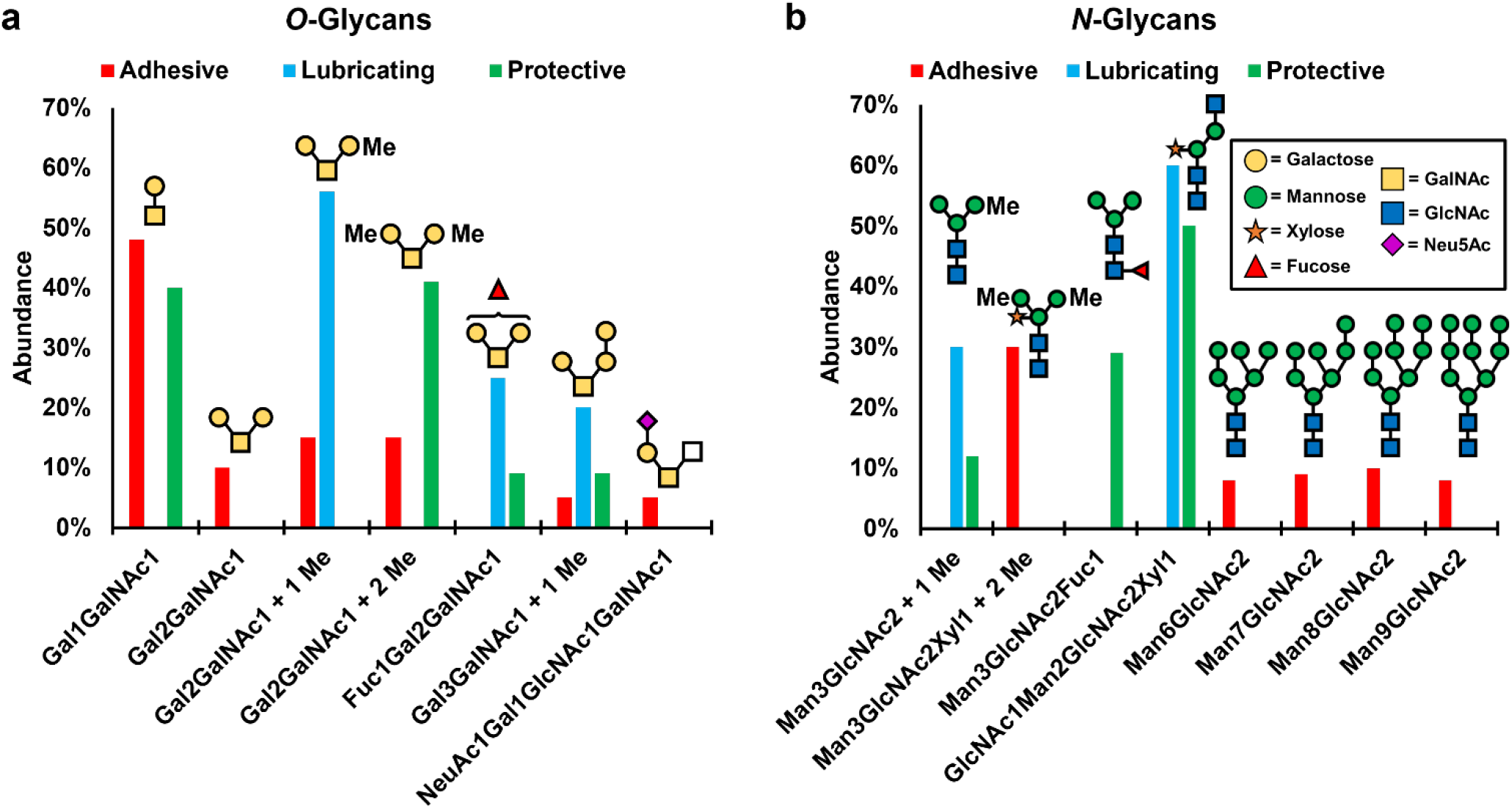
Structures and relative abundance of a) *O*-glycans and b) *N*-glycans found in each isolated mucus secretion. Glycans shown comprised > 5 % of glycomic abundance. Trace *O*- and *N*-glycans (< 5 % abundance) are listed in Supplementary Tables 6 and 7, respectively. Inset in ‘b’ lists monosaccharide structures as defined by the symbolic nomenclature for glycans (SNFG).

Identified glycans are consistent with previous reports on mollusk glycosylation.^57^ In the adhesive and lubricating samples, the mucin core-1 *O*-glycan (T-antigen) was the dominant glycan (52 % and 69 %, respectively). In the protective sample, the most abundant glycan was the trisaccharide (MeGal)_2_GalNAc (48 %), which was the second most-abundant glycan in all other samples. The T-antigen *O*-glycan ((Gal)GalNAc) was the second most abundant glycan in the protective sample. The trisaccharide (Gal)_2_GalNAc and its methylated variants were observed in all three secretions. Adhesive and lubricating mucus contained the unmodified, mono-, and dimethylated versions of this glycan, but the protective mucus only contained the dimethylated form. The preponderance of so few glycans in the samples is surprising, given that human mucin glycans are extremely diverse and typically utilize longer oligosaccharides.^59^ Interestingly, the lubricating mucus showed sizable abundance (∼8 %) of several larger glycans, up to n = 5, while the other mucus possessed only trace amounts of these larger sugars. Gal-rich glycans have a recognized role in biological lubricity,^60^ and increases in polysaccharide length are accompanied with increased material stiffness.^61^ Therefore, longer oligosaccharides likely contribute to its lubricative properties, while the shorter *O*-glycans found in the adhesive and protective mucus would attenuate lubrication. Two sialylated *O*-glycans, Neu5AcGalGlcNAcGalNAc and Neu5AcFuc_2_GalGlcNAcGalNAc, both of which were only found in the adhesive mucus, account for ∼7 % of this sample.

*N-*glycans were extracted from the proteins by treatment with PNGase F and identified by MALDI mass spectrometry (Figure 4b, Supplementary Figures 12–14, Supplementary Table 7).^62^ Twenty-four unique *N*-glycans were detected across all mucus samples. Compositions of these glycans are mainly consistent with *N*-glycans reported in mollusks.^63^ The primary *N-*glycans identified in lubricating mucus are (MeMan)Man_2_GlcNAc_2_ and (Xyl)GlcNAcMan_2_GlcNAc_2_ oligosaccharides.^63^ Protective mucus contained these two structures in addition to significant proportions of Man_3_GlcNAc_2_Fuc and oligomannose sugars. Adhesive mucus contained fifteen unique *N*-linked oligosaccharides not found in the other mucus secretions, with nine containing sialic acids. Six sialylated sugars were found in adhesive mucus, comprising 17 % of the glycomic abundance, while three were observed in the protective mucus, comprising 6 % of the abundance. No sialic acids were detected in the lubricating mucus. Neu5Gc was only detected in adhesive mucus. Interestingly, di- and tri-sialylation was observed exclusively in glycans of the adhesive mucus.

Overall, galactose, GalNAc, mannose, GlcNAc, fucose, xylose, Neu5Ac, and Neu5Gc were identified in the *O*- and *N*-glycans of snail mucus. Fucosylated structures were identified in the *N*-glycans of all three samples, however only the protective and lubricating mucus contained fucosylated *O*-glycans. This observation supports previous accounts of low levels of fucosylation in invertebrate *O*-glycans.^64^ The presence of methylation in both the *N-* and *O*-glycans supports prior reports on both marine and land snails, and identified compositions are similar to those identified in other snails.^65^ A difference is the degree of methylation, as previous studies on other species reported high levels (3 – 4 methyl groups per monosaccharide), while our studies indicate lower levels of methylation (0 – 2 methyl groups).^57^

### Electron microscopy and elemental analysis of *C. aspersum* mucus

Mucus microscale morphologies and elemental composition were determined with SEM and EDX analysis, respectively, on fresh mucus deposited directly onto imaging substrates. The adhesive mucus formed large amorphous masses and ferning patterns that are consistent with mucus secretions in other organisms (Supplementary Figure 15).^66^ Snail lubricious mucus appears oriented into thinner parallel lines. Protective mucus formed sheets that extended across much larger lengths than the other secretions. The elemental analysis of each mucus sample revealed substantial differences (Table 1, Supplementary Figures 16–21). The lubricating mucus contained a higher carbon, oxygen, and nitrogen (organic) content (92.4 %) compared to the adhesive (84.7 %) or protective (82.52 %) secretions. Of particular interest is the amount of Ca^2+^ present in each mucus, as increased CaCO_3_ content has been linked to increased mucus crosslinking and adhesiveness.^3^ Ca^2+^ appears to be present in varying amounts, and was measured to be 0.92 %, 1.93 %, and 3.32 % in the lubricating, adhesive, and protective samples, respectively.

**Table 1.** Elemental composition of the snail mucus identified by energy dispersive X-ray spectroscopy (EDX) analysis. Wt% refers to the percent abundance of each elemental species divided by its atomic molecular mass and normalized.

| **Element** | **Adhesive**<br>**(Wt%)** | **Lubricating**<br>**(Wt%)** | **Protective**<br>**(Wt%)** |
| --- | --- | --- | --- |
| **C** | 58.0 | 79.5 | 62.2 |
| **O** | 15.6 | 8.29 | 18.8 |
| **N** | 9.38 | 4.61 | — |
| **Cl** | 4.50 | 1.15 | 5.87 |
| **K** | 4.11 | 3.00 | 2.55 |
| **Na** | 2.96 | 0.46 | 3.57 |
| **Ca** | 1.93 | 0.92 | 3.32 |
| **S** | 1.80 | — | 1.53 |
| **Mg** | 0.90 | 1.15 | 1.53 |
| **P** | 0.64 | — | 0.26 |
| **Si** | — | 0.92 | 0.38 |

### Measurement of adhesive energy and elastic modulus

Stiffness and adhesion of the mucus hydrogels were characterized with scanning probe analysis. AFM imaging revealed that the adhesive and protective mucus were composed of large aggregates or sheets, while the lubricating mucus contained regions of smaller particles evenly spread across the surface (Supplementary Figure 22). Similar sizes and morphologies were observed for the samples under ambient atmospheric conditions in the AFM and under vacuum in the SEM, suggesting that mucus morphology appears to be resilient to extreme changes in pressure and to desiccation. As such, the cracks observed in the imaging are not likely to be artifacts of the drying process.

AFM nanoindentation spectroscopy determined the mechanical stiffness and energy involved in mucus adherence. As mucus hydrogels are sensitive to moisture conditions, 50 % relative humidity was maintained during the experiments by monitoring the chamber with a humidity sensor and injecting dry or moist air as needed to prevent fluctuations in gel swelling.^67^ Since mucus is a heterogenous material, each sample was subjected to multiple indentations across different regions of the substrate (n = 36 for adhesive, n = 51 for lubricious, and n = 59 for protective) to account for topographical differences, and each indentation produced an approach-retraction force-separation curve (Figure 5a). Curves were fit using the Johnson–Kendall–Roberts (JKR) model,^68^ which accounts for adhesive interactions between the AFM tip and the sample. Curve fittings were then used to calculate Young’s modulus (*E*) and the adhering energy (*W*) for each indentation.^67^ The distributions of *E* (Figure 5b) and *W* (Figure 5c) across all indentations were determined, and average values for each distribution were calculated (Figure 5d). Secreted mucus from other invertebrates have reported *E* of 0.1 – 200 MPa, and all the measured values fall within this range.^69,70^ *E* for the adhesive mucus was significantly lower (41.6 ± 2.03 MPa) than that of the lubricating (132 ± 12.0 MPa) and protective (162 ± 18.6 MPa) samples, indicating adhesive mucus is much less stiff. Mucus from gastropods and other animals have *W* of 2 – 20 N•m^−1^.^71^ *W* for protective mucus (17.2 ± 2.63 N•m^−1^) was much greater than adhesive (3.37 ± 0.498 N•m^−1^) and lubricating (2.39 ± 0.268 N•m^−1^) samples. The greater adhesiveness of protective mucus could increase this secretion’s ability to trap pathogens and other materials.

**Figure 5.**
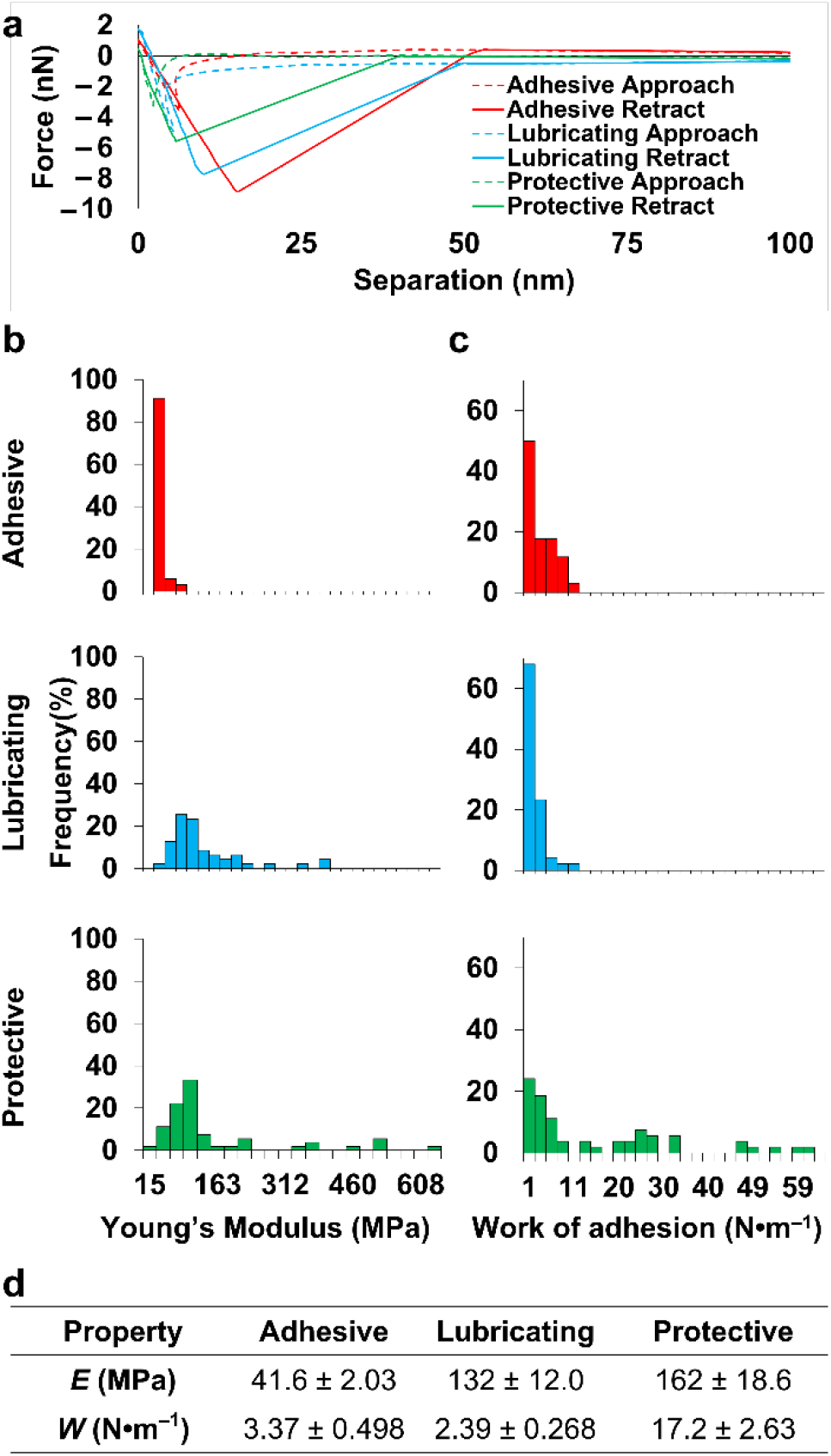
Mechanical properties of adhesive (red), lubricating (blue), and protective (green) snail mucus determined by force-ramp indentation. a) Representative force-distance curve pairs from the adhesive, lubricating, and protective snail mucus. b) Distribution of measured Young’s Modulus, *E*, and c) work of adhesion, *W*, values for each mucus secretion. d) Average values of *E* and *W* for adhesive (n = 36), lubricating (n = 51), and protective (n = 59) mucus from the data plotted in b) and c). Error is defined as standard error, or standard deviation divided by the square root of the number of measurements, n.

## Discussion

Comparative analysis of these datasets revealed the origins of the adhesive, lubricating, and protective behavior of *C. aspersum* mucus (Figure 1b). For example, adhesive mucus has low *E* and high *W* relative to the other secretions, which are desirable properties in biological adhesives.^72^ Adhesive mucus can stretch and self-heal because of the H-bonds, ion bridges, and disulfide linkages that form reversible crosslinks, allowing dynamic sol-gel transitions without sacrificing material properties.^73^ The gel’s ion content also explains its flexibility. Hydrogels containing ionic crosslinks relieve stress through ion bridge reformation (stress relief occurs on a timescale of *t* ∼ 20 s), while covalently crosslinked hydrogels employ water migration to relieve stress (*t* ∼ 1 h), suggesting ion-dependent gels can stretch and spread more readily under mechanical stimuli.^74^ Adhesive mucus contains large amorphous masses, which could increase contact surface area, thereby increasing adhesion between the snail and substrate. Greater glycoprotein,^75^ oligomeric protein,^76^ and glue protein^42^ expression relative to the other mucus also contribute to the material’s adhesiveness. Also, the adhesive mucus exclusively contains tyrosinase, which catalyzes the formation of DOPA, a chemical signature of adhesives secreted by mussels, which is possibly why enzymes involved in its metabolism are found in adhesive mucus.^42^ DOPA-based adhesion has not been reported in snails, and this finding suggests that *C. aspersum* could utilize a similar mechanism as mussels and other mollusks to adhere to inorganic substrates.^42^ Elemental analysis determined this mucus has comparatively high Ca^2+^ content. Secretion of Ca^2+^ likely increases mucus adhesion by coordinating ion bridges in the hydrogel.^3,21^ This idea is further supported by the presence of acidic sialylated glycans, which would increase cation binding within the gel and to substrates. As only the adhesive mucus contains Neu5Gc, it is possible only pedal tissue expresses Neu5Gc-synthesizing enzymes, while the dorsal tissue does not, supported by the fact that Neu5Ac is a biosynthetic precursor of Neu5Gc.^77^ Together, these chemical compositions may generate weak cross-linking that construct a continually reforming flexible adhesive, allowing the snail to adhere to the roughest horizontal, inclined, and inverted surfaces.

*C. aspersum* lubricious mucus was stiff and minimally adhesive, and these properties may provide minimal friction and adhesion on the snail’s pedal surface. Linear structures along its axis of motion reduce surface roughness and, in turn, friction.^78^ A large portion of this hydrogel’s composition (∼40 %) was collagen. Increased collagen levels increase hydrogel stiffness,^79^ which explains the elevated *E* of the lubricious mucus compared to the adhesive (<10 % collagen). Compared to the Ca-rich adhesive mucus, hydrogels formed with crosslinked collagen, like the lubricious mucus, require longer timescales to relieve mechanical stress and thus would have increased *E*.^74^ Ca^2+^-binding proteins were abundant in lubricating mucus, which may sequester free Ca^2+^ and prevent ion bridges from forming. Elemental analysis revealed the gel’s Ca^2+^ content (0.92 %) was less than half of that in the adhesive (1.93 %) and one-third the amount in the protective (3.32 %). Lower salt concentration leads to fewer ionic crosslinks in the gel, resulting in lower *W* of the lubricious mucus.^3^ The elongated oligosaccharides found in the lubricious mucus are extensively hydrated and minimize glycan chain interpenetration under low loads, like those experienced by the snail, which is known to increase lubricity.^55^ The result of all of these elements is a rigid non-stick gel underneath the snail during locomotion, allowing effortless movement across any surface.

*C. aspersum* protective mucus combines features from the adhesive and lubricative mucus, resulting in a hybrid material with the stickiness of the adhesive mucus and the rigidity of lubricious mucus. Protective mucus forms contiguous sheets covering more surface area than the other secretions. Like the adhesive, the protective mucus shows high Ca^2+^ content alongside glue proteins and glycoproteins, thereby potentially increasing *W*. These proteins are modified with short *O*-glycans, which could decrease lubricity. Like the lubricious mucus, the protective mucus has high collagen expression (∼25 % of protein expression), which correlates to high *E*. The stiffness may also be the consequence of ionic, covalent, and noncovalent linkages forming within the hydrogel,^73,74^ which could allow the gel to relieve stress via both ion bridge rearrangement and water migration,^74^ as the role of Ca^2+^ in forming snail mucus ion bridges is well known.^8^ Consequently, mechanical inputs would have their energy dispersed across different timescales. The protective gel would thereby respond to stress more quickly than covalently linked hydrogels but also experience more strain than ionically crosslinked ones. Additionally, these behaviors could indicate a tougher gel that can better maintain its integrity. Though all three mucus contained protease inhibitors, which shield host proteins from degradation,^46^ the protective hydrogel’s distinguishing factor is the presence of lectins, which confer antimicrobial properties to the mucus by preventing pathogenic binding.^49^ This material also possesses diverse *N*-glycans, which likely increase interactions with pathogens in the environment, thereby trapping threats and allowing defensive proteins to engage.

The comparative analysis of the three distinct mucus illuminates the origins of their functional differences (Figure 1b). General principles regarding snail mucus were elucidated, leading to important findings that can be used to advance the field of mucus research. Secreted snail mucus are majorly comprised of collagens and glycoproteins. A relatively simple set of 8 *O*-glycan structures decorate these *C. aspersum* mucus proteins compared to, for example, 76 in *Xenopus laevis*^80^ and 169 in *Salmo salar*.^81^ The *O*-glycan length is modulated between mucus, possibly altering stiffness and lubricity.^55^ Each mucus has drastically different *N*-glycosylation. Sialic acids, which are uncommon in mollusks,^65^ were detected in *O*- and *N*-glycans of the adhesive mucus. Covalent, noncovalent, and ionic crosslinking appears to have a significant effect upon mechanical properties, and snails rely upon defensive proteins to protect hydrogel integrity. Additionally, CAMPs containing *N*-terminal glycodomains and *C*-terminal functional domains were identified, showing there is much to be learned by identifying and annotating mucus genes. The comparative mucomics strategy applied here for *C. aspersum* can be used to determine how compositions of other animal secretions account for their ecological function or to assist in the development of synthetic analogues with similarly advantageous biological and chemical properties.^83,84^

## Experimental Section

### Materials

All chemicals were purchased from VWR unless otherwise noted.

## Methods

### Mucus Collection

Snails were provided in October 2021 by Peconic Escargot (Cutchogue, NY, USA), where they were cultured at room temperature and provided a diet of dirt, wild herbs, and cultivated herbs *ad libitum*. 25 physically active snails that were between 5 – 7 cm were washed with room temperature tap water to remove food, debris, and pathogens and placed into a plastic aquarium. Snails were allowed to crawl freely on petri dishes to collect lubricating mucus. To collect adhesive mucus, snails were placed against an inverted dish until adhered and left suspended for 15 minutes. Lubricating and adhesive mucus were not processed or manipulated further and were immediately place on ice for preservation. Protective mucus secretion was induced by gently rubbing the snail’s back with a spatula, which was scraped off into a plastic test tube. All mucus samples were stored under ice packs without further processing in an insulated cooler for transport to the laboratory, where they were then stored at ‒80 °C until use.

### Mucus Protein Purification

Mucus samples were thawed and physical debris was removed with tweezers. 2 mL of 6 M Guanidinium HCl (Gdn), CsCl (density 1.388 g/mL) was added to mucus-containing petri dishes and incubated at 4 °C overnight to dissolve mucus. Additional residue was collected from the dishes by gently scraping residue with a razor blade. Mucus-containing solutions in the petri dishes were pooled by mucus type into 13.2 mL ultracentrifuge tubes (Beckman-Coulter). Samples were then subjected to isopycnic density gradient ultracentrifugation^85^ in a swinging bucket SW41 Ti rotor ultracentrifuge (35,000 rpm, 72 hr, 4 °C) at a relative centrifugal force of 150,000 x g, within which mucus migrates to a characteristic band and cells are removed from the solution. Following centrifugation, tubes were pierced with a needle and fractionated (0.5 – 1 mL). Each fraction was measured for density and tested for carbohydrate content using a microtiter periodic acid-Schiff’s reagent (PAS) staining protocol.^86^ Fractions with a density of approximately 1.4 g/mL as well as high signal-to-background absorbance at 550 nm, indicating high glycoprotein content, were considered mucus-positive. Mucin-positive fractions were pooled and dithiothreitol (DTT) was added to each pool to reach a final concentration of 0.05 M DTT and shaken at 45 °C overnight in an Echotherm orbital mixing dry bath (Torrey Pines Scientific) to reduce disulfide bonds in the mucus hydrogel networks. Reduced samples were then dialyzed (MM cutoff 2 kDa) against 3 changes of ultrapure water over 48 h and fluffy white precipitate formed. Samples were then lyophilized at ‒55 °C / 1 mbar, resulting in a light beige powder which was stored at ‒80 °C. Protein content was quantified at each step in the purification using a Nanodrop one-C spectrophotometer (Thermo-Fisher).

### RNA Extraction and Sequencing

Snails provided by Peconic Escargot in February 2020 were sacrificed on-site via freezing in a dry ice-ethanol mixture. Whole snails were stored in Invitrogen RNAlater™ (Thermo Fisher, AM7021) and frozen at ‒80 °C until used. 6 individual tissue slices of the snail’s dorsal and pedal surfaces of the foot were excised from different snails. Total RNA was extracted from these slices using a Qiagen RNeasy Micro kit (Qiagen, 74004) according to manufacturer’s instructions. The integrity of total RNA was confirmed using nanodrop and Agilent 2100 BioAnalyzer analysis. The RNA Integrity Number (RIN) was not considered because of known co-migration of 28S rRNA fragments with 18S rRNA in molluscan RNA, causing decreased RIN values in the absence of RNA degradation.^87,88^ Total RNA was used as a template to perform polyA enriched first strand cDNA synthesis using the HiSeq RNA sample preparation kit for Illumina Sequencing (Illumina Inc., CA) following manufacturer’s instructions. The cDNA libraries were sequenced using Illumina HiSeq 1000 technology using a paired end flow cell and 80 × 2 cycle sequencing.

### Read Processing and *De Novo* Assembly

Raw reads were quality checked with FastQC v0.11.5 (www.bioinformatics.babraham.ac.uk).^89^ Adapter sequences and low-quality reads (Phred score <33) were removed using Trimmomatic v0.36 and trimmed reads were re-evaluated with FastQC to ensure the high quality of the data after the trimming process.^90^ Due to the lack of a reference genome, the processed reads were *de novo* assembled using Trinity v2.4.0.^*91*^ *De novo* assembled transcriptomes were translated with Trinity Super Transcripts.^92^ Supertranscripts was used to construct the largest isoform of each gene, in other words producing the original unspliced transcripts, rather than spliced variants of the transcripts.^92^ 179,552 transcripts were assembled. RNA sequences were deposited in Genbank with the primary accession codes SAMN29856567, SAMN29856568, SAMN29856569, SAMN29856570, SAMN29856571, SAMN29856572.

### Proteomic Mass Spectrometry

2 μg of purified snail mucus protein samples at a concentration of 1 mg/mL were loaded onto a single 10% SDS-PAGE stacking mini gel (#4561034, BioRad) band to remove lipids, detergents and salts. The single gel band containing all proteins was reduced with DTT, alkylated with iodoacetic acid and digested with trypsin. 2 μg of extracted peptides were re-solubilized in 0.1% aqueous formic acid and loaded onto a Thermo Acclaim Pepmap (Thermo, 75uM ID X 2cm C18 3uM beads) precolumn and then onto an Acclaim Pepmap Easyspray (Thermo, 75uM X 15cm with 2µM C18 beads) analytical column separation using a Dionex Ultimate 3000 uHPLC at 250 nL/min with a gradient of 2-35% organic (0.1% formic acid in acetonitrile) over 1 hr. Peptides were analyzed using a Thermo Orbitrap Fusion mass spectrometer operating at 120,000 resolution (FWHM in MS1) with HCD sequencing (15,000 resolution) at top speed for all peptides with a charge of 2+ or greater.

### Bioinformatic analysis

The raw data were converted into *.mgf format (Mascot generic format) for searching using the Mascot 2.6.2 search engine (Matrix Science) against predicted sequences from the *de novo* assembled snail transcriptome.^93^ The database search results were loaded onto Scaffold Q+ Scaffold_4.9.0 (Proteome Sciences) for statistical treatment and data visualization.^94^ Peptide identifications were made by exact homology of fragmented peptides against translated transcripts. Using the Scaffold Local FDR (false discovery rate) algorithm, probability thresholds for peptide identifications and protein identifications were set at 95.0% and 5.0%, respectively, to achieve an FDR less than 1.0%, as per proteomic research standards.^18,28^ Additionally, accepted sequences must have contained at least 2 identified peptides. Peptides were quantified by MS/MS counts. Proteomics data were submitted to the PRIDE database under the accession number PXD035534.

The sequences of the proteins identified in the mucus samples were subjected to BLASTP^95^ searches using default parameters to determine their functions based on homology with known proteins in the NCBI non-redundant protein database.^15^ Simultaneously, HMMER^53^ was also used to conduct domain searches against the PFAM^96^ protein database. Employing these two platforms in tandem results in more accurate functional assignments based upon sequence identity and shared domain structures. Each protein was manually classified into one of nine functional categories: lectin, glycoprotein, network-formation, matrix, enzymes, protease inhibitors, ion-binding, regulatory, or housekeeping. Proteins that had sequence similarity with predicted snail proteins without known function were classified as “unknown,” and proteins that had no similarity with any known proteins were classified as “novel.” Using Clustal Omega^97^ within the EMBL–EBI web form (ebi.ac.uk/Tools/msa/clustalo/) using default parameters, a multiple sequence alignment was conducted on our 71 proteins as well as an extensive set of reference proteins to generate a dendrogram and cluster the genes studied via neighbor-joining. For each protein type found, 3 – 5 proteins of the same type from other gastropods or mollusk species were selected from the NCBI non-redundant protein database and added to the alignment. Additionally, other protein types that appeared in the initial BLAST search results were included to build more accurate relationships. By using a global comparative approach,^98^ validation of protein functional assignments and characterization of the more elusive proteins are streamlined. In most cases, proteins of a given type were paired alongside known proteins of the same type, with only minimal cases of orphaned sequences. Molluscan proteins of each functional category, as well as three human mucins, were included in the tree generation. Three proteins of an unrelated family were included as an outgroup. Display and annotation of alignment tree was conducted using iTOL v5.^99^ Sequences were uploaded into the HMMER web server for identification of domains.^53^ Multiple sequence alignment of proteins was conducted using Jalview.^100^

### Release of *N*-glycans

Lyophilized mucin samples were reduced by adding 25 mM DTT and incubating at 45°C for 60 mins. DTT was then removed using Amicon ultracentrifuge 10 kDa spin filters. Samples were then resuspended in 50 mM ammonium bicarbonate and 2 µL of PNGase F (New England Biolabs) was added. Samples were incubated at 37 °C for 48 hrs, and an additional aliquot of PNGase F was added after 24 hrs. Following incubation, released *N*-glycans were separated from the deglycosylated protein by passing through an Amicon ultracentrifuge 10 kDa spin filter. The flow-through was then loaded onto a C18 SPE cartridge (Resprep) and eluted with 5% acetic acid. The *N*-glycan fraction as well as the de-*N*-glycosylated protein fraction were lyophilized.

### Release of *O*-glycans

Dried samples were then dissolved in 0.1 M NaOH and mixed with 55 mg/mL sodium borohydride. Samples were then subjected to 52 hr *β*-elimination at 45°C. Following incubation, samples were neutralized with 10 % acetic acid dropwise and passed through DOWEX H+ resin column and C18 SPE column. Samples were eluted with 5 % acetic acid. Eluted *O*-glycans were then lyophilized. Borates were removed using 9:1 methanol: acetic acid under a stream of N_2_.

### Per-*O*-methylation and Profiling by Matrix-Assisted Laser-Desorption Time-of-Flight Mass Spectrometry (MALDI-TOF-MS)

Dried samples were then dissolved in dimethylsulfoxide (DMSO) and methylated using NaOH/DMSO base and methyl iodide. The reaction was quenched using LC-MS grade water, and Per-*O*-methylated glycans were extracted with methylene chloride and dried under N_2_. Permethylated glycans were dissolved in methanol. Glycans were then mixed 1:1 with α-dihyroxybenzoic acid (DHB) matrix. MALDI-TOF-MS analysis was done in positive ion mode using an AB SCIEX TOF/TOF 5800 mass spectrometer. Glycans were identified according to previously established snail Glycan assignments. Glycomics data were submitted to GlycoPost database under the accession number GPST000297.

### Scanning Electron Microscopy/Energy Dispersive X-ray Spectroscopy

To create the samples, live snails were allowed to crawl on SEM Al pin stubs (Ted Pella, 16144) that were inverted or horizontal, to create samples for adhesive and lubricating mucus, respectively, and back (protective) mucus was scraped onto the stubs, similar to the silicon wafer samples for AFM, and air-dried overnight. The samples were sputter-coated with gold to a thickness of 5 nm using a Leica EM ACE600 Coater for better electrical conductivity. These samples were then imaged in a Thermo Scientific (FEI) Helios NanoLab 660 FIB-SEM with HT of 5 kV, current of 6.3, 13 and 25 pA with ETD (Everhart-Thornley) detector. EDS (energy-dispersive X-ray spectroscopy) mapping was collected with an Oxford detector at HT of 10 kV and current of 1.6 nA. Data was collected and analyzed using AZtec software.^101^

### AFM Topography

To create the samples, live snails were allowed to crawl on Si wafers that were inverted (adhesive) or horizontal (lubricating), while back (protective) was directly deposited onto the wafer and directly analyzed. The samples were subjected to AFM imaging and analysis using an AFM (Multimode 8, Bruker) under ambient temperature (25 °C) and relative humidity (50 %) to mimic conditions experienced by snails in the wild. Mucus topographies were measured by using an AFM probe with a tip radius of ∼2 nm (SCANASYST-AIR, Bruker). All measurements were taken at 50% relative humidity, which was controlled by injecting dry or moist air into the enclosed AFM chamber and measured by a humidity sensor (HIH-4021, Honeywell).

### Stiffness and work of adhesion characterization via the JKR model

The stiffness of mucus samples was characterized using AFM nano-indentation method,^68^ where an indenter (MLCT-E, Bruker) with radius of 20 nm and a spring constant of 0.139 N/m was used. The indentation deflection sensitivity was 40.7 nm/V, calibrated by performing an indentation on the silicon wafer substrate. Peaks of three mucus aggregates are indented to obtain the force *vs*. displacement relationships, of which the retracting portion of the indenting profiles were subsequently analyzed by using the Johnson–Kendall–Roberts (JKR) model, given by

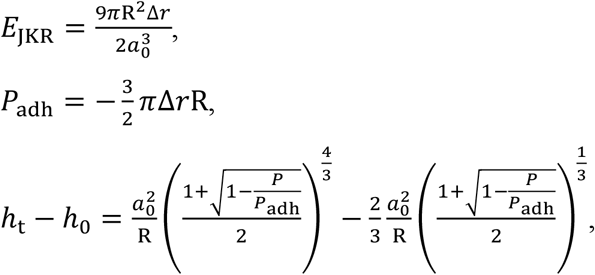

where *E*_JKR_ is the Young’s modulus, R is the tip radius, ∆*r* is the work of adhesion, *a*_0_ is the contact area when the contract force is zero, *P*_adh_ is the pull-off force, *h*_t_ is the indentation depth, *h*_0_ is the contact point where the pull-off force shows, and *P* is the load. The work of adhesion was measured by the area enclosed by the approaching and the retracting indentation force-displacement curves, and was normalized by the probe sample contact area (*a*_0_), given by *a*_0_ = π*Rh*_t_.

## Data availability

Proteomics data that support the findings of this study have been deposited in PRIDE with the primary accession codes [PXD035534]. RNA sequences that support the findings of this study have been deposited in Genbank with the primary accession codes SAMN29856567, SAMN29856568, SAMN29856569, SAMN29856570, SAMN29856571, SAMN29856572. Glycomics data were submitted to GlycoPost database under the accession number GPST000297. The authors declare that all other data that support the findings of this study are available within the paper and its supplementary information files.

## Acknowledgements

A.R.C. acknowledges support from a CUNY Science Scholarship and a CUNY Llewellyn Fellowship. A.B.B. acknowledges support from the Air Force Office of Scientific Research (FA9550-19-1-0220). L.E.P. and P.A. acknowledge the Complex Carbohydrate Research Center (Athens, GA) and support from the National Institutes of Health-funded R24 grant (NIH-R24GM137782) and NSF GlycoMIP, a National Science Foundation Materials Innovation Platform funded through Cooperative Agreement (DMR-1933525). X.C. acknowledges support from the Office of Naval Research (N00014-18-1-2492). A.B.B. and D.B. acknowledge support from the Army Educational Outreach Program (Rochester, NY) and Harlem Educational Activities Fund (New York, NY). MH acknowledges support from Allen Institute’s Distinguished Investigator Award and NIH SPEECH Pilot Project U54CA221704 & U54CA221705. Taylor Knapp and Peconic Escargot (Cutchogue, NY) are acknowledged for providing the animals used in this study. Genevieve Arroyo, Nicholas Mueller, and Robert Gullery are acknowledged for assisting in the collection of snail mucus. The Gardner, Cassacia, and Ulijn labs at the CUNY Advanced Science Research Center are acknowledged for allowing use of lab equipment and space to conduct experiments. NYU’s Genome Technology Center is acknowledged for conducting the transcriptomic sequencing with support from the Laura and Issac Perlmutter Cancer Center (Cancer Center Support Grant P30CA016087). The Clinical Proteomics Platform at the RIMUHC, McGill is acknowledged for conducting the proteomics experiments and analysis.

## Author Contributions

A.B.B. conceived the research. A.B.B., A.R.C, M.H., X.C., L.E.P. and P.A. coordinated the research. A.R.C. and M.B.M. collected the snail mucus samples and snails used in these studies. A.R.C. conducted mucin purifications. L.E.P. extracted glycans and conducted glycomic analysis. A.R.C. and M.B.M. conducted RNA extraction and transcriptomic analysis. Z.-L.L. conducted AFM experiments. A.R.C., Z.-L.L., and D.B. conducted AFM mechanical property analysis. A.R.C. and S.Z. conducted the SEM and EDX experiments. All authors contributed to analysis and discussion of data. The manuscript was written by A.R.C. and A.B.B. and edited and approved by all authors.

## Additional Information

**Supplementary Information** is available for this paper. Reprints and permissions information is available at www.nature.com/reprints.

## References

1 Cerullo, A. R. et al. Comparative animal mucomics: Inspiration for functional materials from ubiquitous and understudied biopolymers. ACS Biomaterials Science & Engineering 6, 5377–5398 (2020).

2 Lieleg, O. & Ribbeck, K. Biological hydrogels as selective diffusion barriers. Trends in Cell Biology 21, 543–551, doi:https://doi.org/10.1016/j.tcb.2011.06.002 (2011).

3 Zhong, T., Min, L., Wang, Z., Zhang, F. & Zuo, B. Controlled self-assembly of glycoprotein complex in snail mucus from lubricating liquid to elastic fiber. RSC advances 8, 13806–13812 (2018).

4 Co, J. Y., Crouzier, T. & Ribbeck, K. Probing the Role of Mucin-Bound Glycans in Bacterial Repulsion by Mucin Coatings. Advanced Materials Interfaces 2, 1500179, doi:https://doi.org/10.1002/admi.201500179 (2015).

5 Allam, B. & Espinosa, E. P. in Mucosal Health in Aquaculture 325–370 (Elsevier, 2015).

6 McShane, A. et al. Mucus. Current Biology 31, R938–R945, doi:https://doi.org/10.1016/j.cub.2021.06.093 (2021).

7 Denny, M. Locomotion: the cost of gastropod crawling. Science 208, 1288–1290 (1980).

8 Hughes, G. W. et al. The MUC5B mucin polymer is dominated by repeating structural motifs and its topology is regulated by calcium and pH. Scientific reports 9, 1–13 (2019).

9 Witten, J., Samad, T. & Ribbeck, K. Molecular Characterization of Mucus Binding. Biomacromolecules 20, 1505–1513, doi:10.1021/acs.biomac.8b01467 (2019).

10 Gabriel, U. I., Mirela, S. & Ionel, J. Quantification of mucoproteins (glycoproteins) from snails mucus, Helix aspersa and Helix Pomatia. Journal of Agroalimentary Processes and Technologies 17, 410–413 (2011).

11 Kimura, K., Chiba, S. & Koene, J. M. Common effect of the mucus transferred during mating in two dart-shooting snail species from different families. Journal of Experimental Biology 217, 1150–1153 (2014).

12 Dolashki, A. et al. Structure and antibacterial activity of isolated peptides from the mucus of garden snail Cornu aspersum. Bulg Chem Commun 50, 195–200 (2018).

13 Vong, A., Ansart, A. & Dahirel, M. Dispersers are more likely to follow mucus trails in the land snail Cornu aspersum. The Science of Nature 106, 43 (2019).

14 Allam, B. & Espinosa, E. P. in Mucosal Health in Aquaculture (eds Benjamin H. Beck & Eric Peatman) 325–370 (Academic Press, 2015).

15 Pruitt, K. D., Tatusova, T. & Maglott, D. R. NCBI Reference Sequence (RefSeq): a curated non-redundant sequence database of genomes, transcripts and proteins. Nucleic acids research 33, D501–D504 (2005).

16 Ballance, S. et al. Partial characterisation of high-molecular weight glycoconjugates in the trail mucus of the freshwater pond snail Lymnaea stagnalis. Comparative Biochemistry and Physiology Part B: Biochemistry and Molecular Biology 137, 475–486 (2004).

17 Yakubov, G. E., Papagiannopoulos, A., Rat, E., Easton, R. L. & Waigh, T. A. Molecular structure and rheological properties of short-side-chain heavily glycosylated porcine stomach mucin. Biomacromolecules 8, 3467–3477 (2007).

18 Ballard, K. R., Klein, A. H., Hayes, R. A., Wang, T. & Cummins, S. F. The protein and volatile components of trail mucus in the Common Garden Snail, Cornu aspersum. PloS one 16, e0251565 (2021).

19 Belouhova, M. et al. Microbial diversity of garden snail mucus. MicrobiologyOpen 11, e1263 (2022).

20 Tachapuripunya, V., Roytrakul, S., Chumnanpuen, P. & E-kobon, T. Unveiling Putative Functions of Mucus Proteins and Their Tryptic Peptides in Seven Gastropod Species Using Comparative Proteomics and Machine Learning-Based Bioinformatics Predictions. Molecules 26, 3475 (2021).

21 Jia, D. & Muthukumar, M. Theory of Charged Gels: Swelling, Elasticity, and Dynamics. Gels 7, 49 (2021).

22 Newar, J. & Ghatak, A. Studies on the adhesive property of snail adhesive mucus. Langmuir 31, 12155–12160 (2015).

23 McDermott, M. et al. Advancing Discovery of Snail Mucins Function and Application. Frontiers in Bioengineering and Biotechnology 9 (2021).

24 Cilia, G. & Fratini, F. Antimicrobial properties of terrestrial snail and slug mucus. Journal of Complementary and Integrative Medicine 15 (2018).

25 Takagi, J. et al. Mucin O-glycans are natural inhibitors of Candida albicans pathogenicity. Nature Chemical Biology 18, 762–773, doi:10.1038/s41589-022-01035-1 (2022).

26 Greistorfer, S. et al. Snail mucus− glandular origin and composition in Helix pomatia. Zoology 122, 126–138 (2017).

27 Sinitcyn, P., Rudolph, J. D. & Cox, J. Computational methods for understanding mass spectrometry–based shotgun proteomics data. Annual Review of Biomedical Data Science 1, 207–234 (2018).

28 Espinosa, E. P., Koller, A. & Allam, B. Proteomic characterization of mucosal secretions in the eastern oyster, Crassostrea virginica. Journal of proteomics 132, 63–76 (2016).

29 Valcourt, U., Alcaraz, L. B., Exposito, J.-Y., Lethias, C. & Bartholin, L. Tenascin-X: beyond the architectural function. Cell Adhesion & Migration 9, 154–165 (2015).

30 Nicholas, B. et al. Shotgun proteomic analysis of human-induced sputum. Proteomics 6, 4390–4401 (2006).

31 Tørresen, O. K. et al. Tandem repeats lead to sequence assembly errors and impose multi-level challenges for genome and protein databases. Nucleic Acids Research 47, 10994–11006, doi:10.1093/nar/gkz841 (2019).

32 Benedito, R. et al. The notch ligands Dll4 and Jagged1 have opposing effects on angiogenesis. Cell 137, 1124–1135 (2009).

33 De Lau, W. B., Snel, B. & Clevers, H. C. The R-spondin protein family. Genome biology 13, 1–10 (2012).

34 Gaur, R. K. Amino acid frequency distribution among eukaryotic proteins. The IIOAB Journal 5, 6 (2014).

35 Lang, T. et al. Searching the Evolutionary Origin of Epithelial Mucus Protein Components-Mucins and FCGBP. Mol Biol Evol 33, 1921–1936, doi:10.1093/molbev/msw066 (2016).

36 Li, D. & Graham, L. D. Epiphragmin, the major protein of epiphragm mucus from the vineyard snail, Cernuella virgata. Comparative Biochemistry and Physiology Part B: Biochemistry and Molecular Biology 148, 192–200 (2007).

37 Mane, P. C. et al. Terrestrial snail-mucus mediated green synthesis of silver nanoparticles and in vitro investigations on their antimicrobial and anticancer activities. Scientific reports 11, 1–16 (2021).

38 Li, Y., Su, J. & Cavaco-Paulo, A. Laccase-catalyzed cross-linking of BSA mediated by tyrosine. International Journal of Biological Macromolecules 166, 798–805 (2021).

39 Park, S.-W. et al. The protein disulfide isomerase AGR2 is essential for production of intestinal mucus. Proceedings of the National Academy of Sciences 106, 6950–6955 (2009).

40 Provan, F. et al. Proteomic analysis of epidermal mucus from sea lice–infected A tlantic salmon, S almo salar L. Journal of fish diseases 36, 311–321 (2013).

41 Chen, Y. et al. FKBP65-dependent peptidyl-prolyl isomerase activity potentiates the lysyl hydroxylase 2-driven collagen cross-link switch. Scientific reports 7, 1–9 (2017).

42 Bilotto, P. et al. Adhesive Properties of Adsorbed Layers of Two Recombinant Mussel Foot Proteins with Different Levels of DOPA and Tyrosine. Langmuir 35, 15481–15490, doi:10.1021/acs.langmuir.9b01730 (2019).

43 Cao, W. et al. Unraveling the Structure and Function of Melanin through Synthesis. Journal of the American Chemical Society 143, 2622–2637, doi:10.1021/jacs.0c12322 (2021).

44 Unlu, A. & Ekici, A. Phenoloxidase is involved in the immune reaction of Helix lucorum to parasitic infestation by dicrocoeliid trematode. Annals of agricultural and environmental medicine: AAEM 28 (2021).

45 Rajan, B. et al. Proteome reference map of the skin mucus of Atlantic cod (Gadus morhua) revealing immune competent molecules. Fish & shellfish immunology 31, 224–231 (2011).

46 Hasnain, S. Z., McGuckin, M. A., Grencis, R. K. & Thornton, D. J. Serine protease (s) secreted by the nematode Trichuris muris degrade the mucus barrier. (2012).

47 Liu, W. et al. Stress-Induced Mucus Secretion and Its Composition by a Combination of Proteomics and Metabolomics of the Jellyfish Aurelia coerulea. Marine Drugs 16, 341 (2018).

48 Allain, T., Fekete, E. & Buret, A. G. Giardia cysteine proteases: the teeth behind the smile. Trends in parasitology 35, 636–648 (2019).

49 Nayak, A., Pednekar, L., Reid, K. B. & Kishore, U. Complement and non-complement activating functions of C1q: a prototypical innate immune molecule. Innate immunity 18, 350–363 (2012).

50 Pietrzyk-Brzezinska, A. J. & Bujacz, A. H-type lectins–Structural characteristics and their applications in diagnostics, analytics and drug delivery. International Journal of Biological Macromolecules 152, 735–747 (2020).

51 Caruana, N. J., Strugnell, J. M., Faou, P., Finn, J. & Cooke, I. R. Comparative proteomic analysis of slime from the striped pyjama squid, Sepioloidea lineolata, and the southern bottletail squid, Sepiadarium austrinum (Cephalopoda: Sepiadariidae). Journal of proteome research 18, 890–899 (2019).

52 Patel, D. M. & Brinchmann, M. F. Skin mucus proteins of lumpsucker (Cyclopterus lumpus). Biochemistry and biophysics reports 9, 217–225 (2017).

53 Potter, S. C. et al. HMMER web server: 2018 update. Nucleic acids research 46, W200–W204 (2018).

54 Moniaux, N., Escande, F., Porchet, N., Aubert, J.-P. & Batra, S. K. Structural organization and classification of the human mucin genes. Front Biosci 6, D1192–D1206 (2001).

55 Crouzier, T. et al. Modulating Mucin Hydration and Lubrication by Deglycosylation and Polyethylene Glycol Binding. Advanced Materials Interfaces 2, 1500308 (2015).

56 Wilkinson, H. & Saldova, R. Current methods for the characterization of O-glycans. Journal of Proteome Research 19, 3890–3905 (2020).

57 Stepan, H. et al. O-Glycosylation of snails. Glycoconjugate journal 29, 189–198 (2012).

58 Kang, P., Mechref, Y., Kyselova, Z., Goetz, J. A. & Novotny, M. V. Comparative glycomic mapping through quantitative permethylation and stable-isotope labeling. Analytical chemistry 79, 6064–6073 (2007).

59 Jin, C. et al. Structural diversity of human gastric mucin glycans. Molecular & Cellular Proteomics 16, 743–758 (2017).

60 Jay, G. D. & Waller, K. A. The biology of lubricin: near frictionless joint motion. Matrix Biology 39, 17–24 (2014).

61 Wheeler, R. et al. Bacterial cell enlargement requires control of cell wall stiffness mediated by peptidoglycan hydrolases. MBio 6, e00660–00615 (2015).

62 Valk-Weeber, R. L., Dijkhuizen, L. & van Leeuwen, S. S. Large-scale quantitative isolation of pure protein N-linked glycans. Carbohydrate research 479, 13–22 (2019).

63 Staudacher, E. Mollusc N-glycosylation: Structures, Functions and Perspectives. Biomolecules 11, 1820 (2021).

64 Thomès, L. & Bojar, D. The Role of Fucose-Containing Glycan Motifs Across Taxonomic Kingdoms. Frontiers in Molecular Biosciences 8, doi:10.3389/fmolb.2021.755577 (2021).

65 Staudacher, E. Mucin-Type O-Glycosylation in Invertebrates. Molecules (Basel, Switzerland) 20, 10622–10640, doi:10.3390/molecules200610622 (2015).

66 Masmali, A. M., Purslow, C. & Murphy, P. J. The tear ferning test: a simple clinical technique to evaluate the ocular tear film. Clinical and Experimental Optometry 97, 399–406 (2014).

67 Danielsen, S. P. O. et al. Molecular Characterization of Polymer Networks. Chemical Reviews 121, 5042–5092, doi:10.1021/acs.chemrev.0c01304 (2021).

68 Wu, G., Gotthardt, M. & Gollasch, M. Assessment of nanoindentation in stiffness measurement of soft biomaterials: kidney, liver, spleen and uterus. Scientific reports 10, 1–11 (2020).

69 Fudge, D. S., Gardner, K. H., Forsyth, V. T., Riekel, C. & Gosline, J. M. The mechanical properties of hydrated intermediate filaments: insights from hagfish slime threads. Biophysical journal 85, 2015–2027 (2003).

70 Wilks, A. M., Rabice, S. R., Garbacz, H. S., Harro, C. C. & Smith, A. M. Double-network gels and the toughness of terrestrial slug glue. Journal of Experimental Biology 218, 3128–3137, doi:10.1242/jeb.128991 (2015).

71 Newar, J., Verma, S. & Ghatak, A. Effect of Metals on Underwater Adhesion of Gastropod Adhesive Mucus. ACS omega 6, 15580–15589 (2021).

72 Dastjerdi, A. K., Pagano, M., Kaartinen, M., McKee, M. & Barthelat, F. Cohesive behavior of soft biological adhesives: experiments and modeling. Acta Biomaterialia 8, 3349–3359 (2012).

73 Picchioni, F. & Muljana, H. Hydrogels based on dynamic covalent and non covalent bonds: a chemistry perspective. Gels 4, 21 (2018).

74 Zhao, X., Huebsch, N., Mooney, D. J. & Suo, Z. Stress-relaxation behavior in gels with ionic and covalent crosslinks. Journal of applied physics 107, 063509 (2010).

75 Opell, B. D. & Stellwagen, S. D. Properties of orb weaving spider glycoprotein glue change during Argiope trifasciata web construction. Scientific reports 9, 1–11 (2019).

76 Forsprecher, J., Wang, Z., Goldberg, H. A. & Kaartinen, M. T. Transglutaminase-mediated oligomerization promotes osteoblast adhesive properties of osteopontin and bone sialoprotein. Cell adhesion & migration 5, 65–72 (2011).

77 Li, Y. & Chen, X. Sialic acid metabolism and sialyltransferases: natural functions and applications. Applied microbiology and biotechnology 94, 887–905 (2012).

78 Figueroa-Morales, N., Dominguez-Rubio, L., Ott, T. L. & Aranson, I. S. Mechanical shear controls bacterial penetration in mucus. Scientific reports 9, 1–10 (2019).

79 Pokki, J., Zisi, I., Schulman, E., Indana, D. & Chaudhuri, O. Magnetic probe-based microrheology reveals local softening and stiffening of 3D collagen matrices by fibroblasts. Biomedical Microdevices 23, 27, doi:10.1007/s10544-021-00547-2 (2021).

80 Guerardel, Y. et al. O-glycan variability of egg-jelly mucins from Xenopus laevis: characterization of four phenotypes that differ by the terminal glycosylation of their mucins. Biochemical Journal 352, 449–463 (2000).

81 Benktander, J. et al. Effects of Size and Geographical Origin on Atlantic salmon, Salmo salar, Mucin O-Glycan Repertoire*[S]. Molecular & cellular proteomics 18, 1183–1196 (2019).

82 Tailford, L. E., Crost, E. H., Kavanaugh, D. & Juge, N. Mucin glycan foraging in the human gut microbiome. Frontiers in genetics 6, 81 (2015).

83 Kwan, C.-S., Cerullo, A. R. & Braunschweig, A. B. Design and Synthesis of Mucin-Inspired Glycopolymers. ChemPlusChem 85, 2704–2721 (2020).

84 Lema, M. A. et al. Scalable Preparation of Synthetic Mucins via Nucleophilic Ring-Opening Polymerization of Glycosylated N-Carboxyanhydrides. Macromolecules 55, 4710–4720, doi:10.1021/acs.macromol.1c02477 (2022).

85 Corfield, A. P. Glycoprotein methods and protocols: The mucins. Vol. 125 (Springer Science & Business Media, 2000).

86 Kilcoyne, M., Gerlach, J. Q., Farrell, M. P., Bhavanandan, V. P. & Joshi, L. Periodic acid–Schiff’s reagent assay for carbohydrates in a microtiter plate format. Analytical biochemistry 416, 18–26 (2011).

87 Dheilly, N. M. et al. A family of variable immunoglobulin and lectin domain containing molecules in the snail Biomphalaria glabrata. Developmental & Comparative Immunology 48, 234–243, doi:https://doi.org/10.1016/j.dci.2014.10.009 (2015).

88 Barcia, R., Lopez-García, J. M. & Ramos-Martínez, J. I. The 28S fraction of rRNA in molluscs displays electrophoretic behaviour different from that of mammal cells. IUBMB Life 42, 1089–1092 (1997).

89 Brown, J., Pirrung, M. & McCue, L. A. FQC Dashboard: integrates FastQC results into a web-based, interactive, and extensible FASTQ quality control tool. Bioinformatics 33, 3137–3139 (2017).

90 Bolger, A. M., Lohse, M. & Usadel, B. Trimmomatic: a flexible trimmer for Illumina sequence data. Bioinformatics 30, 2114–2120 (2014).

91 Haas, B. J. et al. De novo transcript sequence reconstruction from RNA-seq using the Trinity platform for reference generation and analysis. Nature protocols 8, 1494–1512 (2013).

92 Davidson, N. M., Hawkins, A. D. & Oshlack, A. SuperTranscripts: a data driven reference for analysis and visualisation of transcriptomes. Genome biology 18, 1–10 (2017).

93 Helsens, K., Martens, L., Vandekerckhove, J. & Gevaert, K. MascotDatfile: an open-source library to fully parse and analyse MASCOT MS/MS search results. Proteomics 7, 364–366 (2007).

94 Searle, B. C. Scaffold: a bioinformatic tool for validating MS/MS-based proteomic studies. Proteomics 10, 1265–1269 (2010).

95 Johnson, M. et al. NCBI BLAST: a better web interface. Nucleic acids research 36, W5–W9 (2008).

96 El-Gebali, S. et al. The Pfam protein families database in 2019. Nucleic acids research 47, D427–D432 (2019).

97 Thompson, J. D., Gibson, T. J. & Higgins, D. G. Multiple sequence alignment using ClustalW and ClustalX. Current protocols in bioinformatics, 2.3. 1-2.3. 22 (2003).

98 Duraisamy, S., Ramasamy, S., Kharbanda, S. & Kufe, D. Distinct evolution of the human carcinoma-associated transmembrane mucins, MUC1, MUC4 AND MUC16. Gene 373, 28–34 (2006).

99 Letunic, I. & Bork, P. Interactive Tree Of Life (iTOL) v5: an online tool for phylogenetic tree display and annotation. Nucleic acids research 49, W293–W296 (2021).

100 Procter, J. B. et al. in Multiple Sequence Alignment 203–224 (Springer, 2021).

101 Burgess, S. & Pinard, P. AZtec Wave–a New Way to Achieve Combined EDS and WDS Capability on SEM. Microscopy and Microanalysis 26, 114–115 (2020).

