## Supporting Information for "Comparative Mucomic Analysis of Three Functionally Distinct *Cornu aspersum* Secretions"

**Table of Contents**

1. Mucus collection……………………………………………………………….……..S3
2. Mucus purification……………………………………………………………….....S4
3. Proteomic analysis……….…………………………………………………….....S9
4. Glycomic Analysis…..………………………………...………..……………..S21
5. Scanning electron microscopy………………………………………………….S31
6. Atomic force microscopy analysis……………………..………………………….S39

### 1. Mucus collection

**General methods.** In October 2021, snail mucus was collected directly from *C. aspersum* snails (provided by Peconic Escargot in Cutchogue, NY) in three separate manners to differentiate by function, as described previously.[1,2] Snails were cultured at room temperature and provided a diet of dirt, wild herbs, and cultivated herbs *ad libitum*. 25 physically active snails that were between 5 – 7 cm were washed with room temperature tap water to remove food, debris, and pathogens and placed into a plastic aquarium. Lubricating mucus was collected by allowing snails to crawl along horizontal petri dishes to deposit the secretion used to facilitate movement. Adhesive mucus was collected by holding snails against inverted petri dishes to induce adhesion, and the snails were left suspended for 15 min such that they deposited onto the dish the mucus used in adhesion. Lubricating and adhesive mucus were not processed or manipulated further and were immediately place on ice for preservation. Protective mucus was collected by gently scraping the dorsal surface of the snail with a spatula, collecting the skin mucus and depositing it in a collection tube. Samples were stored under ice packs without furhter processing in an insulated cooler for transport and then stored in an ultralow temperature (–80 ˚C) freezer until use.

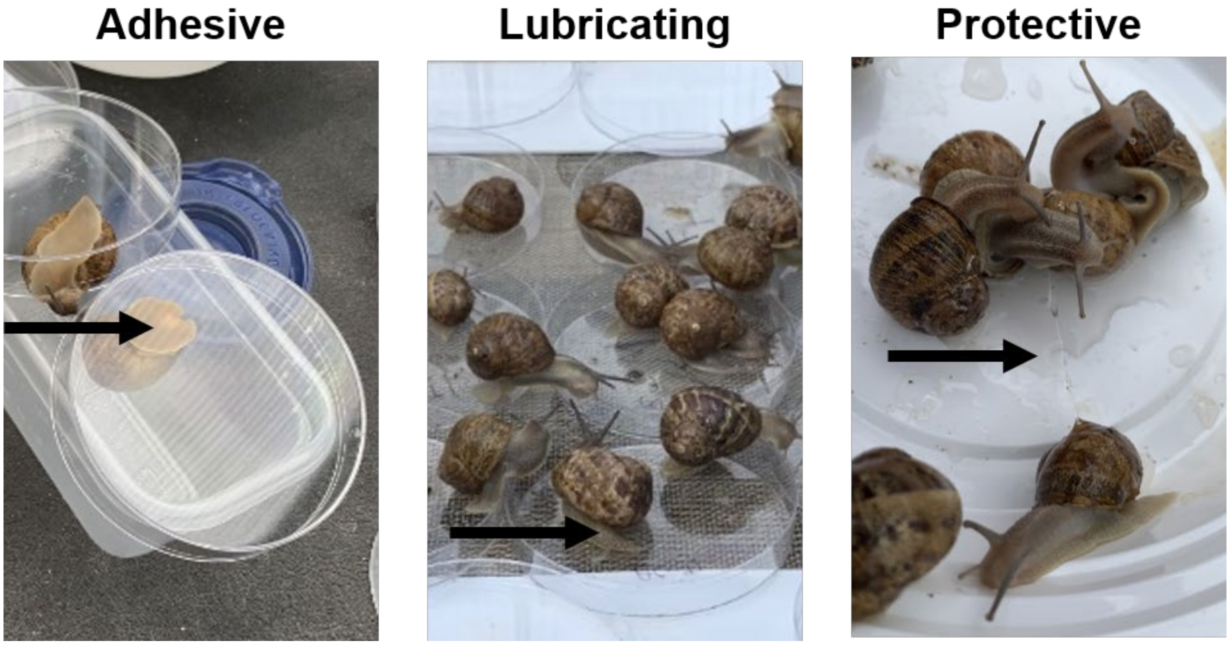


**Figure S1.** Collection of mucus from *C. aspersum* snails. Arrows point to secreted mucus.

### 2. Mucus purification.

**General methods.** Mucus samples were thawed and physical debris was removed with tweezers. 2 mL of 6 M Guanidinium HCl (Gdn), CsCl (density 1.388 g/mL, measured by gravimetric analysis) was added to mucus-containing petri dishes and collection tubes and incubated at 4 ºC overnight on an Ohaus RockingShaker orbital shaker to dissolve mucus, as described previously.[3] After overnight incubation, the mucus-containing solutions in the petri dishes were pooled by mucus type into 13.2 mL ultracentrifuge tubes (Beckman-Coulter). Additional Gdn solution and mucus residue were collected by gently scraping the petri dishes with a razor blade. Samples were then subjected to isopycnic density gradient ultracentrifugation in a swinging bucket SW41 Ti rotor in a Beckman-Coulter Optima XE Ultracentrifuge (35,000 rpm, 72 hr, 4 ºC), at a relative centrifugal force of 150,000 x g, within which mucus migrates to a characteristic band and cells would be removed from the solution, as described previously.[4] Following centrifugation, tubes were pierced with a needle and fractionated (0.5 – 1 mL fractions). Notably, the mucus samples were colorless aggregates about two-thirds up the gradient. Additionally, each fraction was measured for density and tested for carbohydrate content using a microtiter periodic acid-Schiff's reagent (PAS) staining protocol, as described previously.[5] 25 µL of each fraction was added to each well of a clear, flat-bottom 96 well plate. 120 µL of 0.06 % w/v perioidic acid, 7 % v/v glacial acetic acid in water, was added to each well in the dark and covered in tin foil and left to incubate for 90 min at 25 ˚C. After incubation, 100 µL of Schiff's reagent was added to each well in the dark and covered to incubate for 60 min at 25 ˚C. The plate was then subjected to spectrophotometric analysis, and absorbance was measured at 550 nm using a Molecular Devices SpectraMAX 190 microplate reader. Additionally, the densities of each fraction were determined by gravimetric analysis, measuring the mass of 500 µL of each fraction. Fractions with a density of approximately 1.4 g/mL as well as high signal-to-background absorbance at 550 nm were considered mucin-positive because it has been reported mucins exhibit a characteristic buoyancy, migrating to this fraction of the density gradient, and glycans labelled with Schiff's stain exhibit an absorbance maximum at 550 nm.[4] Mucin-positive fractions were pooled and dithiothreitol (DTT) was added to each pool to reach a final concentration of 0.05 M DTT, and shaken at 45 ºC overnight in an Echotherm orbital mixing dry bath (Torrey Pines Scientific) to reduce disulfide bonds in the mucus hydrogel networks. Reduced samples were then dialyzed in a cellulose membrane (MM cutoff 2 kDa) against 3 changes of ultrapure water over 48 h and flocculent beige precipitate formed. Samples were then lyophilized using a Labconco Freezedry-System / Freezone 4.5 at –55 ºC / 1 mbar, resulting in a light beige powder which was stored at –80 ºC. Protein content in each collected mucus sample was quantified at each step in the purification using the Nanodrop one-C spectrophotometer (Thermo-Fisher), comparing values to protein concentration standard curves. Standard curves of absorbance vs. protein concentration (in mg/mL) were generated using the same Nanodrop one-C spectrophotometer, and tracking the absorbance signal at 280 nm for varying concentrations of protein. Solutions for the standard curve were generated by dissolving dried mucus protein in 6 M Gdn solution and mixing.

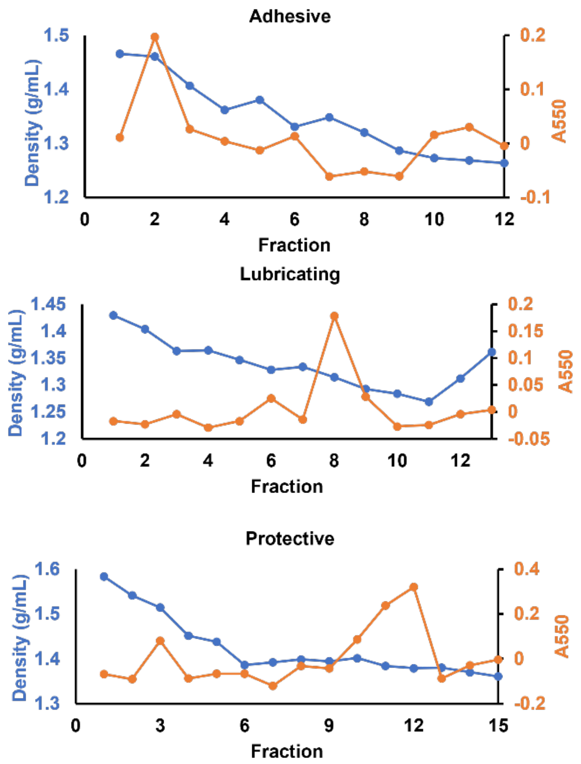


**Figure S2.** Fractionation of snail mucins through isopycnic density gradient ultracentrifugation. Blue curves represent measured density of each fraction. Orange curves represent periodic acid-Schiff's stain (PAS) response using a previously established microtiter plate assay.[5]

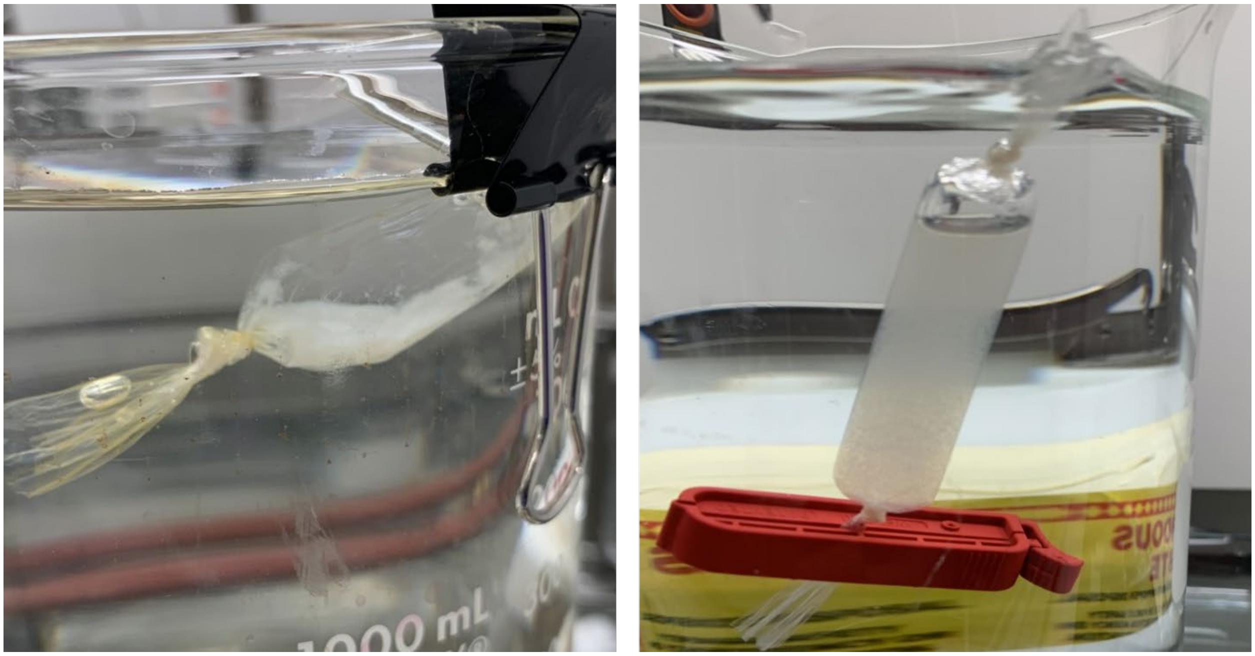

**Figure S3.** Mucus samples post-dialysis. Flocculent beige precipitate forms after dialyzing mucin against ultrapure water from Guandinium hydrochloride solution. Left image is after allowing sample to sit undisturbed for at least 30 min so that precipitate sediments. Right image is immediately after mixing, dispersing protein evenly throughout the suspension. Solutions were clear and colorless prior to dialysis.

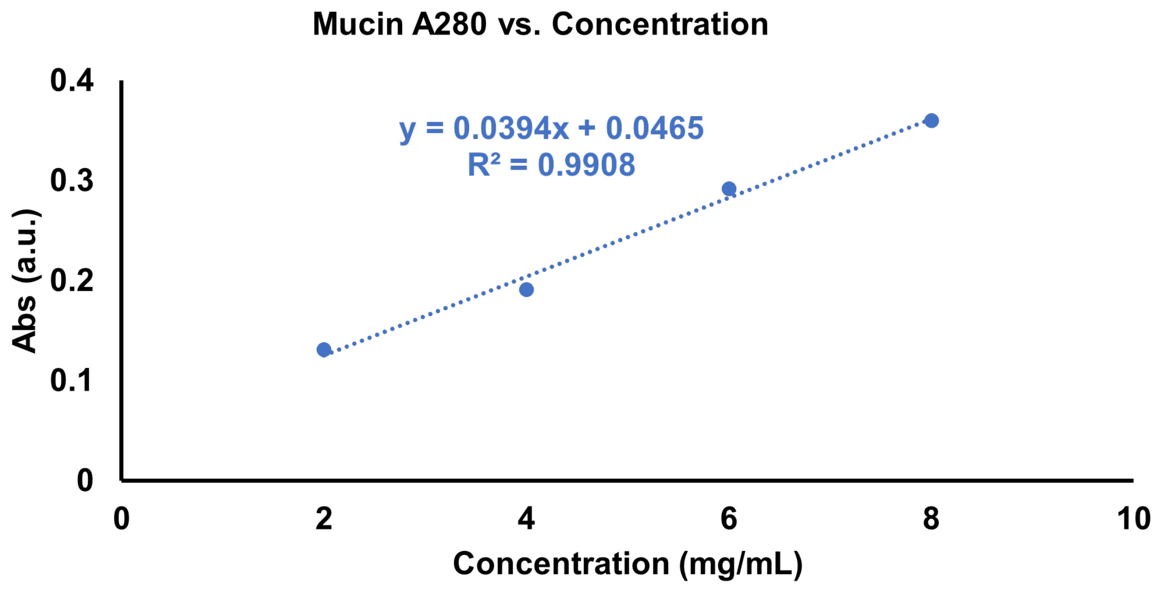


**Figure S4.** Mucus standard curve for both purified mucus proteins determined by Nanodrop spectrophotometric analysis using a linear fit.

**Table S1.** Concentrations in mg/mL were calculated according to mucus protein standard curve in Supplementary Figure 4. UC stands for ultracentrifugation.

| Sample | Protein Content pre-UC (mg) | Protein Content post-UC (mg) | Protein Content post-Dialysis (mg) | % of initial mass |
|---|---|---|---|---|
| Adhesive | 77.2 | 24.6 | 2.98 | 3.86 |
| Lubricating | 168.0 | 29.2 | 12.4 | 7.38 |
| Protective | 120.0 | 30.0 | 5.60 | 4.67 |

### 3. Proteomic analysis

*RNA Extraction and Sequencing*

Snails provided by Peconic Escargot in February 2020 were sacrificed on-site via freezing in a dry ice-ethanol mixture. Whole snails were stored in Invitrogen RNAlater™ (Thermo Fisher, AM7021) and frozen at –80 °C until used. 6 individual tissue slices of the snail's dorsal and pedal surfaces of the foot were excised from different snails. Total RNA was extracted from these slices using a Qiagen RNeasy Micro kit (Qiagen, 74004) according to manufacturer's instructions. The integrity of total RNA was confirmed using nanodrop and Agilent 2100 BioAnalyzer analysis. . The RNA Integrity Number (RIN) was not considered because of known co-migration of 28S rRNA fragments with 18S rRNA in molluscan RNA.[6,7] Total RNA was used as a template to perform polyA enriched first strand cDNA synthesis using the HiSeq RNA sample preparation kit for Illumina Sequencing (Illumina Inc., CA) following manufacturer's instructions. The cDNA libraries were sequenced using Illumina HiSeq 1000 technology using a paired end flow cell and 80 x 2 cycle sequencing.

*Read Processing and De Novo Assembly*

Raw reads were quality checked with FastQC v0.11.5 (www.bioinformatics.babraham.ac.uk).[8] Adapter sequences and low-quality reads (Phred score <33) were removed using Trimmomatic v0.36 and trimmed reads were re-evaluated with FastQC to ensure the high quality of the data after the trimming process.[9] Due to the lack of a reference genome, the processed reads were de novo assembled using Trinity v2.4.0.[10] De novo assembled transcriptomes were translated with Trinity Super Transcripts.[11] Supertranscripts was used to construct the largest isoform of each gene, in other words producing the original unspliced transcripts, rather than spliced variants of the transcripts.[11] 179,552 transcripts were assembled. RNA sequences were deposited in Genbank with the primary accession codes SAMN29856567, SAMN29856568, SAMN29856569, SAMN29856570, SAMN29856571, SAMN29856572.

*Proteomic Mass Spectrometry*

Purified snail mucus protein samples were loaded onto a single stacking gel band to remove lipids, detergents and salts. The single gel band containing all proteins was reduced with dithiothreitol (DTT), alkylated with iodoacetic acid and digested with trypsin. 2 µg of extracted peptides were re-solubilized in 0.1 % aqueous formic acid and loaded onto a Thermo Acclaim Pepmap (Thermo, 75 µM ID X 2 cm C18 3 µM beads) precolumn and then onto an Acclaim Pepmap Easyspray (Thermo, 75 µM X 15 cm with 2 µM C18 beads) analytical column separation using a Dionex Ultimate 3000 uHPLC at 250 nL/min with a gradient of 2-35 % organic (0.1 % formic acid in acetonitrile) over 1 hr. Peptides were analyzed using a Thermo Orbitrap Fusion mass spectrometer operating at 120,000 resolution (FWHM in MS1) with HCD sequencing (15,000 resolution) at top speed for all peptides with a charge of 2+ or greater.

*Proteomic Data Processing*

The raw data were converted into *.mgf format (Mascot generic format) for searching using the Mascot 2.6.2 search engine (Matrix Science) against predicted sequences from the *de novo* assembled snail transcriptome.[12] The database search results were loaded onto Scaffold Q+ Scaffold_4.9.0 (Proteome Sciences) for statistical treatment and data visualization.[13] Peptide identifications were made by exact homology of fragmented peptides against translated transcripts. Using the Scaffold Local FDR (false discovery rate) algorithm, probability thresholds for peptide identifications and protein identifications were set at 95.0 % and 5.0 %, respectively, to achieve an FDR less than 1.0 %, as per proteomic research standards.[14-16] Additionally, accepted sequences must have contained at least 2 identified peptides. Peptides were quantified by MS/MS counts. Proteomics data were submitted to the PRIDE database under the accession number PXD035534.

*Bioinformatic Analysis*

The sequences of the proteins identified in the mucus samples were subjected to BLASTP searches using default parameters to determine their functions based on homology with known proteins in the NCBI non-redundant protein database.[17] Each protein was manually classified into one of nine functional categories: lectin, glycoprotein, network-formation, matrix, enzymes, protease inhibitors, ion-binding, regulatory, or housekeeping. Proteins that had similarity with predicted snail proteins without known function were classified as "Snail," and proteins that had no similarity with any known proteins were classified as "Novel." Sequences were uploaded into ClustalW to generate a dendrogram.[18] Molluscan proteins of each functional category, as well as three human mucins, were included in the tree generation. Protein sequences from *Amphioctopus fangsiao, Aplysia californica, Argopecten irradians, Biomphalaria glabrata, Bulinus truncates, Cernuella virgata, Cornu aspersum, Crassostrea gigas, Crassostrea hongkongensis, Crassostrea virginica, Elysia marginata, Gigantopelta aegis, Haliotis discus, Haliotis rubra, Haliotis tuberculata, Helix pomatia, Hemitoma cumingii, Homo sapiens, Mercenaria mercenaria, Meretrix meretrix, Mizuhopecten yessoensis, Mus musculus, Mytilus coruscus, Mytilus edulis, Mytilus galloprovincialis, Octopus sinensis, Onchidium reevesii, Patella vulgata, Pectin Maximus, Pinctada fucata, Plakobranchus ocellatus, Pomacea canaliculata, Sepia pharaonis,* and *Vampyroteuthis infernalis,* were used to generate the dendrogram. Three proteins from *Mus musculus*, (Pikachurin1, Pikachurin2, and Pikachurin3), were included as an outgroup. Display and annotation of dendrogram was conducted using iTOL v5.[19] Sequences were uploaded into the HMMER web server for identification of domains.[20] Multiple sequence alignment of proteins was conducted using Jalview.[21]

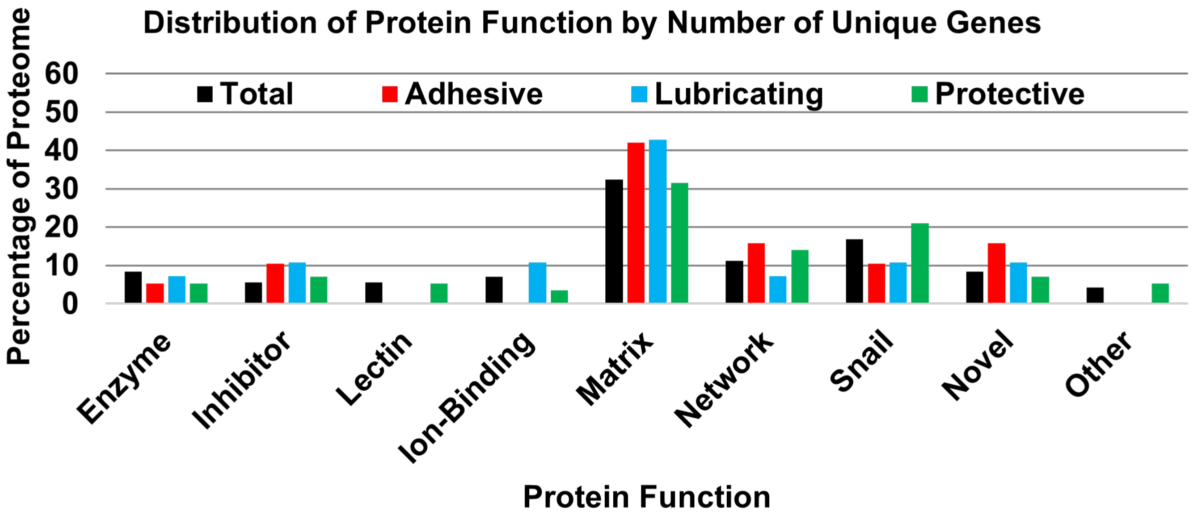


**Figure S5.** Percentage of unique genes for protein function found. Percentages were calculated by finding the ratio of the number of unique genes for each function to the total number of genes within each mucus sample. "Snail" refers to proteins without any determinable function but had structural similarity to uncharacterized proteins previously found in snails. "Novel" indicates the protein had no similarity to any known proteins in the NCBI nor PFAM databases.

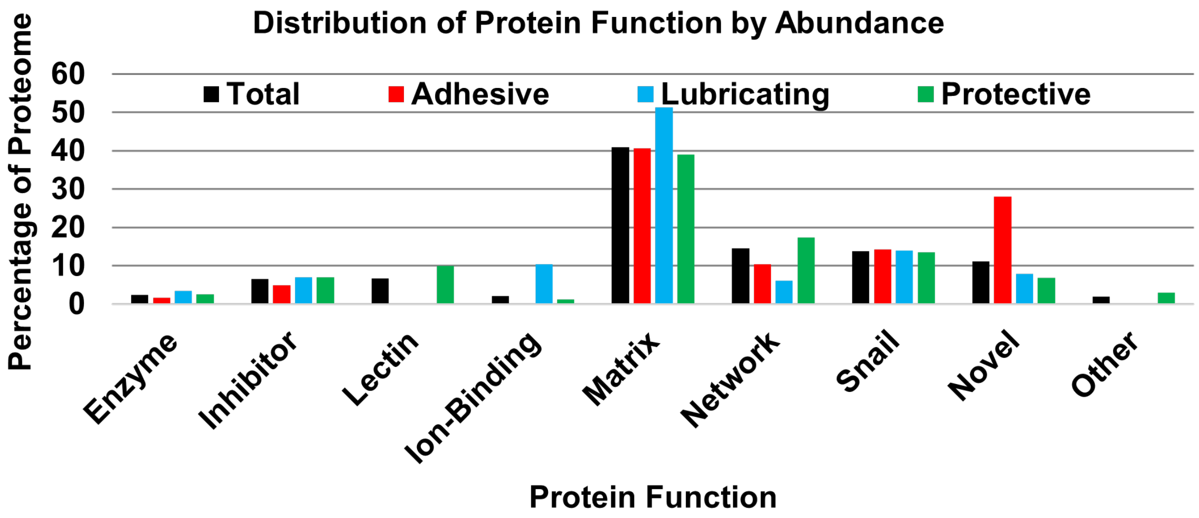


**Figure S6**. Percentage of protein abundance for each function found. Percentages were calculated by finding the ratio of the number of MS/MS counts for all genes within each function to the total number of MS/MS counts within each mucus sample. "Snail" refers to proteins without any determinable function, but had structural similarity to uncharacterized proteins previously found in snails. "Novel" indicates the protein had no similarity to any known proteins in the NCBI nor PFAM databases.

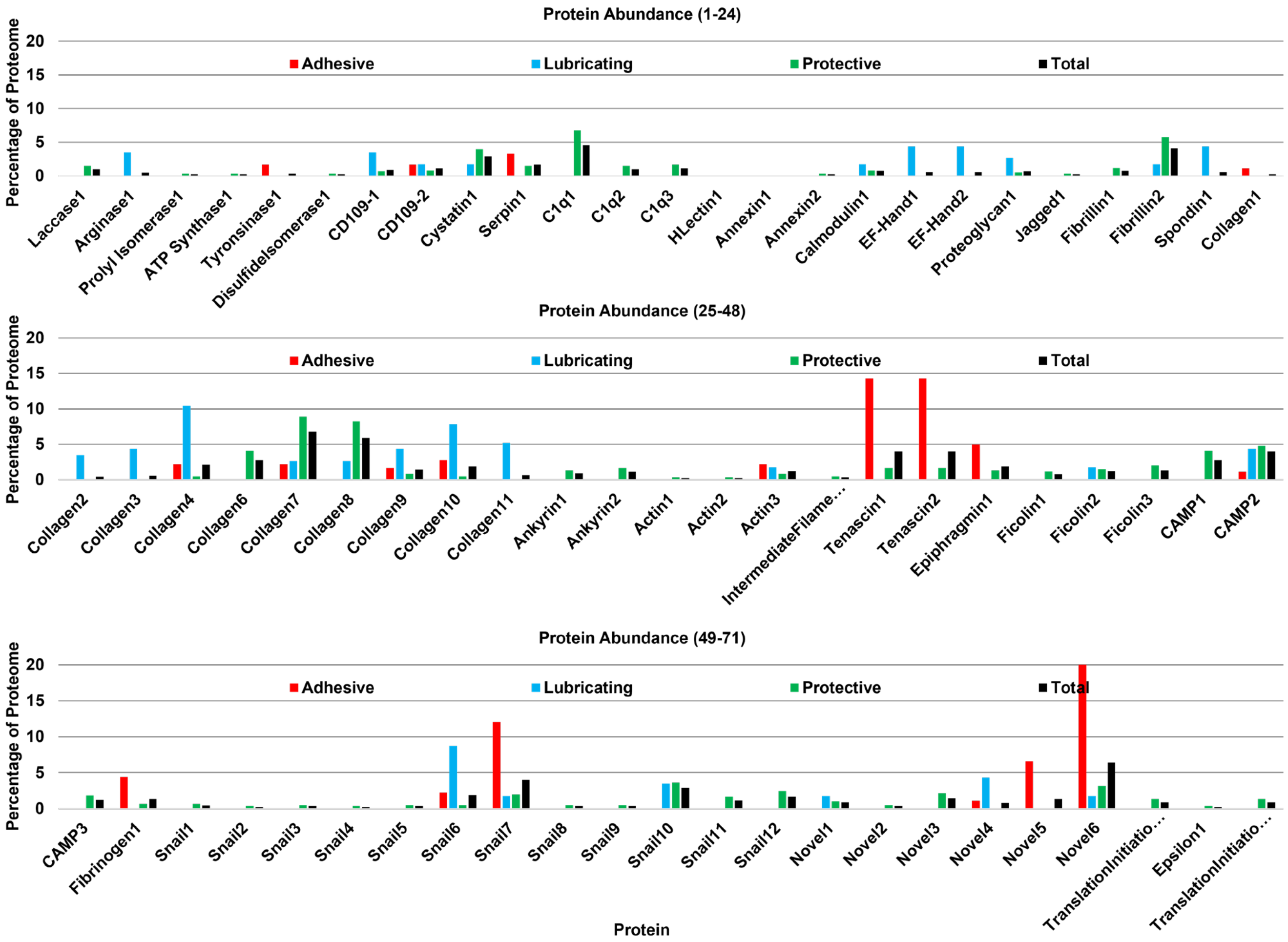


**Figure S7**. Abundances for all 71 proteins identified. Percentages were calculated by finding the ratio of the number of MS/MS counts for each gene to the total number of MS/MS counts within each mucus sample. “Snail” refers to proteins without any determinable function but had structural similarity to uncharacterized proteins previously found in snails. “Novel” indicates the protein had no similarity to any known proteins in the NCBI nor PFAM databases.

| Entry | Protein Name | Accession Number | *MW* | Dendrogram Category | Adhesive Counts | Lubricating Counts | Protective Counts |
|---|---|---|---|---|---|---|---|
| 1 | Actin1 | MM4_TRINITY_DN22220_c0_g1_1 | 40 kDa | Actins | | | 2 |
| 2 | Actin2 | MM4_TRINITY_DN22220_c0_g3_1 | 15 kDa | Actins | | | 2 |
| 3 | Actin3 | MM6_TRINITY_DN19769_c0_g1_1 | 167 kDa | Actins | 4 | 2 | 5 |
| 4 | Ankyrin1 | MM4_TRINITY_DN21122_c3_g1_1 | 54 kDa | Glycoproteins | | | 8 |
| 5 | Ankyrin2 | MM3_TRINITY_DN18245_c0_g1_1 | 34 kDa | Lectins | | | 10 |
| 6 | Annexin1 | MM1_TRINITY_DN12628_c0_g1_1 | 7 kDa | Gel-forming Mucins | | | |
| 7 | Annexin2 | MM3_TRINITY_DN8126_c0_g1_1 | 14 kDa | Annexins | | | 2 |
| 8 | Arginase1 | MM4_TRINITY_DN16115_c0_g1_1 | 32 kDa | Arginases | | 4 | |
| 9 | ATP-Synthase1 | MM6_TRINITY_DN18259_c0_g2_1 | 58 kDa | ATP Synthases | | | 2 |
| 10 | CAMP1 | MM1_TRINITY_DN11035_c0_g1_1 | 46 kDa | Gel-forming Mucins | | | 25 |
| 11 | CAMP2 | MM3_TRINITY_DN18384_c0_g2_1 | 48 kDa | Lectins | 2 | 5 | 29 |
| 12 | CAMP3 | MM4_TRINITY_DN22211_c1_g4_1 | 10 kDa | Gel-forming Mucins | | | 11 |
| 13 | C1q1 | MM5_TRINITY_DN15947_c0_g1_1 | 22 kDa | C1qs | | | 41 |
| 14 | C1q2 | MM4_TRINITY_DN21887_c0_g1_1 | 84 kDa | Lectins | | | 9 |
| 15 | C1q3 | MM5_TRINITY_DN18999_c2_g1_1 | 79 kDa | Lectins | | | 10 |
| 16 | Calmodulin1 | MM4_TRINITY_DN21173_c6_g2_1 | 79 kDa | Ca-binding Proteins | | 2 | 5 |
| 17 | HLectin1 | MM3_TRINITY_DN18322_c4_g2_1 | 28 kDa | Lectins | | | |
| 18 | CD109-1 | MM4_TRINITY_DN25046_c0_g1_1 | 10 kDa | CD-109s | | 4 | 4 |
| 19 | CD109-2 | MM6_TRINITY_DN13823_c0_g1_1 | 53 kDa | CD-109s | 3 | 2 | 5 |
| 20 | Collagen1 | MM3_TRINITY_DN8024_c0_g1_1 | 7 kDa | Matrix Proteins | 2 | | |
| 21 | Collagen2 | MM3_TRINITY_DN17949_c3_g1_1 | 37 kDa | Gel-forming Mucins | | 4 | |
| 22 | Collagen3 | MM4_TRINITY_DN21282_c0_g2_1 | 55 kDa | Gel-forming Mucins | | 5 | |
| 23 | Collagen4 | MM4_TRINITY_DN21282_c0_g5_1 | 31 kDa | Matrix Proteins | 4 | 12 | 3 |
| 24 | Collagen5 | MM5_TRINITY_DN17478_c0_g1_1 | 65 kDa | Collagens | | | 25 |
| 25 | Collagen6 | MM6_TRINITY_DN18929_c0_g3_1 | 30 kDa | Collagens | 5 | 9 | 3 |
| 26 | Collagen7 | MM5_TRINITY_DN19063_c0_g1_1 | 90 kDa | Collagens | 4 | 3 | 54 |
| 27 | Collagen8 | MM5_TRINITY_DN19063_c0_g2_1 | 74 kDa | Collagens | | 3 | 50 |
| 28 | Collagen9 | MM6_TRINITY_DN18929_c0_g1_1 | 57 kDa | Collagens | 3 | 5 | 5 |
| 29 | Collagen10 | MM4_TRINITY_DN17262_c0_g1_1 | 18 kDa | Gel-forming Mucins | | 6 | |
| 30 | Cystatin1 | MM5_TRINITY_DN17535_c0_g1_1 | 23 kDa | Cystatins | | 2 | 24 |
| 31 | DisulfideIsomerase1 | MM1_TRINITY_DN11276_c5_g2_1 | 14 kDa | Gel-forming Mucins | | | 2 |
| 32 | EF-Hand1 | MM4_TRINITY_DN21829_c0_g1_1 | 44 kDa | Ca-binding Proteins | | 5 | |
| 33 | EF-Hand2 | MM6_TRINITY_DN19244_c4_g2_1 | 37 kDa | Ca-binding Proteins | | 5 | |
| 34 | ElongationFactor1 | MM4_TRINITY_DN20761_c0_g1_1 | 62 kDa | Unclustered | | | 5 |
| 35 | Epiphragmin1 | MM6_TRINITY_DN20583_c0_g1_1 | 7 kDa | Gel-forming Mucins | 9 | | 8 |
| 36 | Epsilon1 | MM5_TRINITY_DN5476_c0_g1_1 | 11 kDa | Unclustered | | | 2 |
| 37 | Fibrillin1 | MM3_TRINITY_DN13586_c0_g1_1 | 42 kDa | Glycoproteins | | | 7 |
| 38 | Fibrillin2 | MM3_TRINITY_DN17952_c1_g1_1 | 49 kDa | Matrix Proteins | | 2 | 35 |
| 39 | Fibrinogen1 | MM1_TRINITY_DN10983_c2_g2_1 | 10 kDa | Gel-forming Mucins | 8 | | 4 |
| 40 | Ficolin1 | MM1_TRINITY_DN11240_c0_g1_1 | 41 kDa | Unclustered | | | 7 |
| 41 | Ficolin2 | MM3_TRINITY_DN18425_c0_g1_1 | 66 kDa | Unclustered | | 2 | 9 |
| 42 | IntermediateFilament1 | MM4_TRINITY_DN28955_c0_g1_1 | 11 kDa | Gel-forming Mucins | | | 3 |
| 43 | Jagged1 | MM2_TRINITY_DN9611_c0_g1_1 | 31 kDa | Glycoproteins | | | 2 |
| 44 | Laccase1 | MM3_TRINITY_DN15913_c0_g1_1 | 54 kDa | Laccases | | | 9 |
| 45 | Novel2 | MM3_TRINITY_DN17597_c0_g1_1 | 24 kDa | Lectins | | | 3 |
| 46 | Novel3 | MM3_TRINITY_DN18249_c5_g4_1 | 64 kDa | Glycoproteins | | | 13 |
| 47 | Novel4 | MM4_TRINITY_DN21224_c1_g1_1 | 9 kDa | Matrix Proteins | 2 | 5 | |
| 48 | Novel5 | MM4_TRINITY_DN21844_c0_g1_1 | 14 kDa | Gel-forming Mucins | 12 | | |
| 49 | Novel6 | MM6_TRINITY_DN21806_c0_g1_1 | 9 kDa | Matrix Proteins | 37 | 2 | 19 |
| 50 | ProlylIsomerase1 | MM6_TRINITY_DN18216_c0_g1_1 | 63 kDa | Prolyl Isomers | | | 2 |
| 51 | Proteoglycan1 | MM1_TRINITY_DN9148_c0_g1_1 | 32 kDa | Glycoproteins | | 3 | 3 |
| 52 | Serpin1 | MM5_TRINITY_DN18792_c2_g1_1 | 77 kDa | Serpins | 6 | | 9 |
| 53 | Snail1 | MM1_TRINITY_DN11259_c0_g2_1 | 67 kDa | Glycoproteins | | | 4 |
| 54 | Snail2 | MM1_TRINITY_DN10906_c0_g2_1 | 30 kDa | Lectins | | | 2 |
| 55 | Snail3 | MM2_TRINITY_DN9959_c0_g1_1 | 40 kDa | Lectins | | | 3 |
| 56 | Snail4 | MM5_TRINITY_DN16950_c0_g1_1 | 18 kDa | Mucins | 4 | 10 | 3 |
| 57 | Snail5 | MM5_TRINITY_DN18004_c6_g1_1 | 77 kDa | Glycoproteins | 22 | 2 | 12 |
| 58 | Snail6 | MM4_TRINITY_DN19690_c0_g1_1 | 43 kDa | Gel-forming Mucins | | | 2 |
| 59 | Snail7 | MM4_TRINITY_DN21671_c4_g6_1 | 25 kDa | Glycoproteins | | | 3 |
| 60 | Snail8 | MM6_TRINITY_DN19636_c3_g1_1 | 70 kDa | Lectins | | 4 | 22 |
| 61 | Snail9 | MM6_TRINITY_DN19640_c2_g1_1 | 112 kDa | Glycoproteins | | | 15 |
| 62 | Snail10 | MM4_TRINITY_DN21618_c2_g4_1 | 29 kDa | Lectins | | | 10 |
| 63 | Snail11 | MM5_TRINITY_DN19011_c0_g2_1 | 62 kDa | Lectins | | | 3 |
| 64 | Snail12 | MM6_TRINITY_DN19602_c1_g1_1 | 51 kDa | Lectins | | | 3 |
| 65 | Spondin1 | MM4_TRINITY_DN481_c0_g1_1 | 64 kDa | Glycoproteins | | 5 | |
| 66 | Tenascin1 | MM3_TRINITY_DN18052_c0_g1_1 | 76 kDa | Glycoproteins | 26 | | 10 |
| 67 | Tenascin2 | MM2_TRINITY_DN10241_c6_g1_1 | 76 kDa | Glycoproteins | 26 | | 10 |
| 68 | Ficolin3 | MM4_TRINITY_DN22280_c4_g1_1 | 87 kDa | Glycoproteins | | | 12 |
| 69 | Novel1 | MM3_TRINITY_DN18425_c0_g2_1 | 14 kDa | CD-109s | | 2 | 6 |
| 70 | TranslationInitiationFactor1 | MM1_TRINITY_DN11217_c5_g1_1 | 9 kDa | Unclustered | | | 8 |
| 71 | Tyrosinase1 | MM6_TRINITY_DN19765_c0_g2_1 | 72 kDa | Tyrosinases | 3 | | |

**Table S2**. Proteins identified in proteomic analysis. Quantification of proteins are based on counts in MS/MS analysis. Corresponding transcriptome accession numbers shown.

| Protein | Group | Function |
|---|---|---|
| Laccase | Enzyme | Catalyzes oxidation of phenolic compounds |
| Arginase | | Catalyzes formation of urea from arginine |
| Prolyl Isomerase | | Interconverts cis-trans isomers of proline peptide bonds |
| ATP Synthase | | Produces ATP from ADP |
| Tyrosinase | | Catalzyes formation of DOPA from tyrosine and melanins |
| Disulfide Isomerase | | Catalyzes formation and breakage of disulfide bonds |
| CD109 | Inhibitor | Serine protease inhibitor |
| Cystatin | | Cysteine protease inhibitor |
| Serpin | | Serine protease inhibitor |
| C1q | Lectin | Lectin, binds with serine proteases |
| H-Type Lectin | | GalNAc-binding lectin |
| Annexin | Ion-Binding | Calcium ion regulation |
| Calmodulin | | Calcium-binding messenger protein |
| EF-Hand | | Calcium-binding signaling protein |
| Proteoglycan | Matrix | Heavily glycosylated protein found in the extracellular matrix |
| Jagged-1 | | Cell-surface signalling protein |
| Fibrillin | | Secreted glycoprotein that forms elastic fibers |
| Spondin | | Secreted glycoprotein involved in extracellular matrix |
| Collagen | | Main structural protein of the extracellular matrix |
| Actin | | Cytoskeleton structural protein |
| Tenascin | | Extracellular matrix glycoprotein |
| Epiphragmin | Network | Main protein component of snail epiphragm (adhesive seal) |
| Ficolin | | Oligomeric lectins with collagen- and fibrinogen-like domains |
| Mucin | | Main glycoprotein component of mucus |
| Fibrinogen | | Glycoproteins that form oligomeric networks |

**Table S3**. Specific proteins found in *C. aspersum* snail mucus and their known functions. Snail, Novel, and Other categories not included.

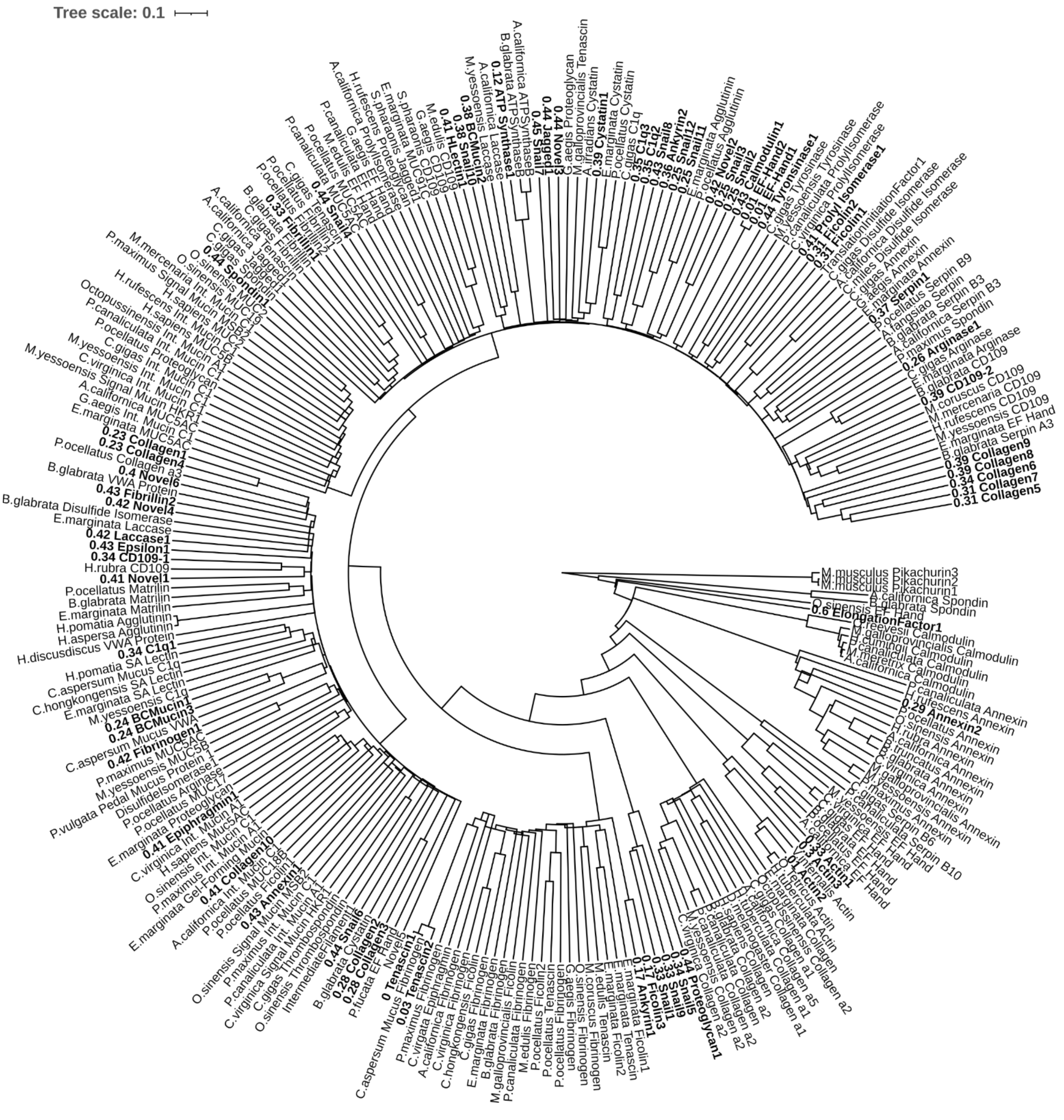


**Figure S8.** Dendrogram of identified snail mucus proteins based on sequence similarity. Topology is identical to the dendrogram shown in Figure 2. The 71 proteins identified in this study are bolded with branch lengths shown. Protein sequences from *Amphioctopus fangsiao, Aplysia californica, Argopecten irradians, Biomphalaria glabrata, Bulinus truncates, Cernuella virgata, Cornu aspersum, Crassostrea gigas, Crassostrea hongkongensis, Crassostrea virginica, Elysia marginata, Gigantopelta aegis, Haliotis discus, Haliotis rubra, Haliotis tuberculata, Helix pomatia, Hemitoma cumingii, Mercenaria mercenaria, Meretrix meretrix, Mizuhopecten*

*yessoensis, Mus musculus, Mytilus coruscus, Mytilus edulis, Mytilus galloprovincialis, Octopus sinensis, Onchidium reevesii, Patella vulgata, Pectin Maximus, Pinctada fucata, Plakobranchus ocellatus, Pomacea canaliculata, Sepia pharaonis,* and *Vampyroteuthis infernalis,* were used to generate the dendrogram.

| Protein | Clade | Putative Function |
|---|---|---|
| Snail1 | Mollusk Glycoproteins | Proteoglycan |
| Snail2 | Lectins | Agglutinin |
| Snail3 | Lectins | Agglutinin |
| Snail4 | Mollusk Mucins | Mucin |
| Snail5 | Mollusk Glycoproteins | Proteoglycan |
| Snail6 | Gel-Forming Mucins | Mucin |
| Snail7 | Mollusk Glycoproteins | Jagged-1 |
| Snail8 | Lectins | Lectin |
| Snail9 | Mollusk Glycoproteins | Proteoglycan |
| Snail10 | Lectins | H-Lectin |
| Snail11 | Lectins | Lectin |
| Snail12 | Lectins | Lectin |
| Novel1 | CD109s | CD109 |
| Novel2 | Lectins | Agglutinin |
| Novel3 | Mollusk Glycoproteins | Proteoglycan |
| Novel4 | Matrix Proteins | Fibrillin |
| Novel5 | Gel-Forming Mucins | Mucin |
| Novel6 | Matrix Proteins | Collagen |

**Table S4**. Proposed functions for uncharacterized proteins based on phylogenetic analysis. “Snail” refers to proteins without any determinable function but had structural similarity to uncharacterized proteins previously found in snails. “Novel” indicates the protein had no similarity to any known proteins in the NCBI nor PFAM databases.

| Protein | Sequence |
|---|---|
| **CAMP1** | NYLRFRIGISGFICGVLLVVVSVTTIGQGERIVFNAKPTTISPGVT<br>PELTVRCGLEDDGNSGVSRVNSIIIRTVDGSVQKEVARIAYRQAAT<br>GGFSTEGASVTGDLSNKAGYLQITWPSPRHGLAGQYNCDIAALATV<br>GDIVKFKSSIRVVSTGKIADLSLSPAFWSAQVKMMAVQTRSNANAQ<br>KTLRHRKRLGTVKGNLILRKRMIRRISAAVQFTTKRALFRSEILIL<br>GENHKLTKRALFRAEILILGENKKKGRRSKSASQLPMPGTRCWSFL<br>PWFWMRIIQESSDTELESILRQEPAGGHFSRGSGCGYKKPQIQNWN<br>QGCHTHGFQGCLWHLVGHNPVSYCPVGQCGITLSQPQTYTCAIIDS<br>NNRHQSGFWLVSLPYVFVHQYQMYVPTCIVIFSTIQLWFLSSKX |
| **CAMP2** | HLSHTDTQYMCVLGSMKEDSISIPIHGCGVFYLQTDRTIVGLSVVC<br>CLLLSLQQVKVKESYSTQSQRLSLQQVKVKESSSRQHQQPSPREHQ<br>SFVFSATPITISPGVTPELTVRCGLEDDGNSGVSRVNSIIIRTVDG<br>SVQKEVARIAYRQAATGGFSTEGASVTGDLSNKAGYLQITWPSPRH<br>GLAGQYNCDIAALATVGDIVKFKSSIRVVSTGKIADLSLSPAFWSA<br>QVKMMAVQTRSNANAQKTLRHRKRLGTVKGNLILRKRMIRRISAAV<br>QFTTKRALFRAAILILGENKKIRDSDSWGEPLKARETVVEVRFSTS<br>YARNPLLVVSPVVLDADNTTPGTRWWSFLPWFWMWLIQKTSDTELE<br>SSDTELESKIIRYRIRIRDVTPTGFKVVCGTWWDTILYRIDVRWVS<br>NVARCLSHRHTHVQS |
| **CAMP3** | GDLSDKAGYLQVTWPSPEHGLAGQYTCDIVAVAESGDNIKFKSSIQ<br>VVSTGKNADSSDSKQCQCTTDIEALKKAVRDSQGKFDSLEKTVNDL<br>KTSX |

**Table S5**. Amino acid sequences of CAMPs.

### 4. Glycomic analysis

*Reduction and N-glycan Release*

Following purification, the lyophilized samples were resuspended in 25 μL of 50 mM $NH_4HCO_3$ buffer. To this, 25 μL of 25 mM dithiothreitol (DTT) was added and the samples were vortexed. The samples were then incubated at 50°C for 45 minutes. Following incubation, the samples were allowed to come to room temperature. Samples were then cleaned and desalted using Amicon Ultra 10 kDa molecular weight cut off (MWCO) filters (Millipore). The filters were first filled with water (500 μL) and centrifuged at 14 000 x g for 10 mins. Resultant flow through was discarded. Then, the sample mixture was loaded onto the filter, and the filter was again centrifuged for 10 min at 14 000 x g. 500 μL of 50 mM $NH_4HCO_3$ was then loaded onto the filter and centrifuged one more time at 14 000 x g for 10 minutes. The flow through was discarded, and the desalted mucin sample remained in the filter. To remove the sample from the filter, the filter was inverted into a clean tube and centrifuged for 1 minute. The filter was then rinsed with 20 μL of 50 mM $NH_4HCO_3$, inverted into the tube containing the sample, and again centrifuged for 1 minute. To this, 2 μL of PNGase F (New England Biolabs) was added. The samples were briefly vortexed and then incubated at 37°C for 48 hours.

Following incubation with PNGase F, the samples were once again passed through a 10 kDa MWCO filter using the conditions stated above. Following centrifugation, 500 μL of 50 mM ammonium bicarbonate was added and centrifuged once more. The flow through, which contained the released *N*-glycans, was then loaded onto a C18 SPE cartridge (Resprep). The C18 cartridge was first washed with 1 mL of methanol (MeOH) and conditioned with 3 mL of 5% acetic acid. The samples were then loaded onto the C18 column and allowed to flow through. N-glycans were then eluted from the column with 3 mL of 5% acetic acid, and the resultant flow through was lyophilized. The de-N-glycosylated protein sample, which remains in the filter, was removed from the filter as stated above. The de-*N*-glycosylated samples were then lyophilized.

*β-elimination and O-glycan Release*

Following lyophilization, the de-*N*-glycosylated protein was subjected to *β*-elimination. The samples were dissolved in 250 μL of 100 mM NaOH solution and vortexed. The pH of the sample was then checked using pH paper to ensure basic conditions (pH ranged from 10-13). Then, 55 mg/mL of $NaBH_4$ in 100 mM NaOH was added and the sample was vortexed. The sample was then incubated at 50°C for 52 hours. Following incubation samples were neutralized by adding 10% acetic acid dropwise, and vortexed between each addition. As the acid is added the sample bubbles, and this process was repeated until bubbling ceased.

Poly-prep chromatography columns (Bio-Rad) were packed with $H^+$ activated ion exchange resins (DOWEX$^{TM}$ 50W x 8-100) and rinsed 5 times with 1 mL of 5% acetic acid. Following rinsing, the neutralized samples were loaded onto the DOWEX columns and allowed to flow through. The flow through was then loaded onto a C18 SPE column, which was washed and conditioned as described above. 3mL of 5% acetic acid was then loaded onto the DOWEX column to elute the oligosaccharides, and this flow through was once again loaded onto the C18 SPE cartridge. Samples were then frozen on dry ice and lyophilized.

Following lyophilization the borates were removed using 9:1 MeOH: acetic acid. 500 μL of the mixture was added to the samples, vortexed and dried under a stream of $N_2$ gas. This process was repeated until the borates were fully removed (approximately 5 times).

*Per-O-methylation of N- and O-linked Glycans*

The *N*- and *O*-linked glycans were then per-*O*-methylated using NaOH/dimethyl sulfoxide (DMSO) base and iodomethane. First, the NaOH/DMSO base was made by adding 100 µL of 50% (v/v) NaOH to a clean, dry glass vial. Then 200 µL of MeOH was added and the solution was then mixed. 4 mL of DMSO was then added and the vial was mixed vigorously for 3 minutes. The sample was then centrifuged at 3000 rcf for 5 mins. A white precipitate forms at the top of the vial, and a clear base at the bottom. The white precipitate and all remaining DMSO were removed without disturbing the base, and 4 mL more of DMSO was added. This procedure was repeated until white precipitate no longer forms (approximately 3 times). 1 mL of DMSO was then added and homogenized with the clear base.

The samples were dissolved in 200 µL of DMSO and vortexed. Then, 300 µL of the NaOH/DMSO base was added, and to this, 100 µL of iodomethane, or iodomethane-D3 was added and the sample was mixed vigorously for 20 minutes using a shaker.[22] The reaction was then quenched using 2 mL of LC-MS grade water. Turbidity was and should be observed. The iodomethane was then removed by bubbling $N_2$ into the sample for approximately 3 minutes. Then, 2 mL of dichloromethane was added, and the sample was mixed vigorously. The sample was then centrifuged at 3000 rcf for 1 minute for phase separation. The upper water layer was then removed, and to additional mL of water was added. This process was repeated a total of five times. Following the last wash, all traces of water was removed, and the dichloromethane fraction was transferred to a clean glass vial. The sample was then dried under a stream of $N_2$. The sample was then resuspended in 20 µL of MeOH and analyzed using MALDI-TOF-MS.

*Mass Spectrometry of Per-O-methylated glycans*

2 µL of the per-*O*-methylated samples were mixed with 2 µL of 2,5-dihydroxybenzoic acid (DHB) MALDI matrix. DHB was made at a concentration of 15 mg/mL in 70:30:0.1 acetonitrile: water: formic acid. 1 µL of the mixture was spotted on a MALDI plate and allowed to dry. The MALDI plate (stainless steel) was then analyzed using an AB Sciex TOF/TOF 5800 System Mass Spectrometer in positive ion mode using reflector mode. This system contains a nitrogen laser (337 nm wavelength) and the laser intensity used was 6400 (arbitrary units). Only MS1 data was collected, no MS/MS or in-source decay was performed. 2500 shots per spectrum were collected.

*Quantification of Glycans using Internal Standard*

To determine the amount of glycan in the samples and internal standard of a known concentration was added. Xylotetraose (Megazyme) was permethylated separately using the procedure outlined above. 0.1 µg of the xylose standard was added to the oligosaccharide-DHB mixture and spotted on a MALDI plate. Concentrations of the glycans released from the mucin samples were determined by comparing the peak intensities to that of the standard.

*Assignment of Glycan Structures*

Glycan compositions were determined using SCIEX Data Explorer software,[23] GlycoWorkbench 2.0 and manual interpretation.[24] Structural assignments were determined based on mass measurement, literature and biological probability. Glycomics data were submitted to GlycoPost database under the accession number GPST000297.

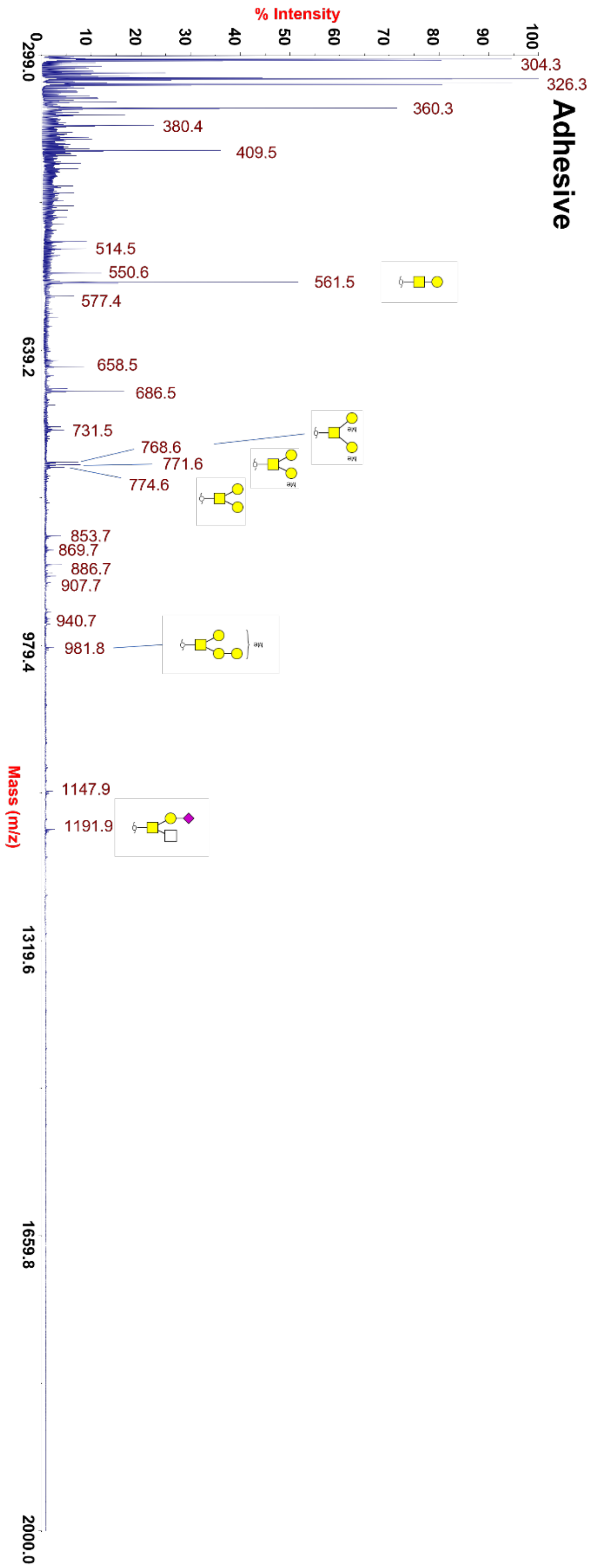


**Figure S9.** Glycomic mass spectrum showing m/z peaks and compositional assignments of extracted *O*-glycans from *C. aspersum* adhesive mucus samples. Insets show magnified region where low-intensity glycan peaks were found. Samples were per-deuter-*O*-methylated and adducts shown are Na+.

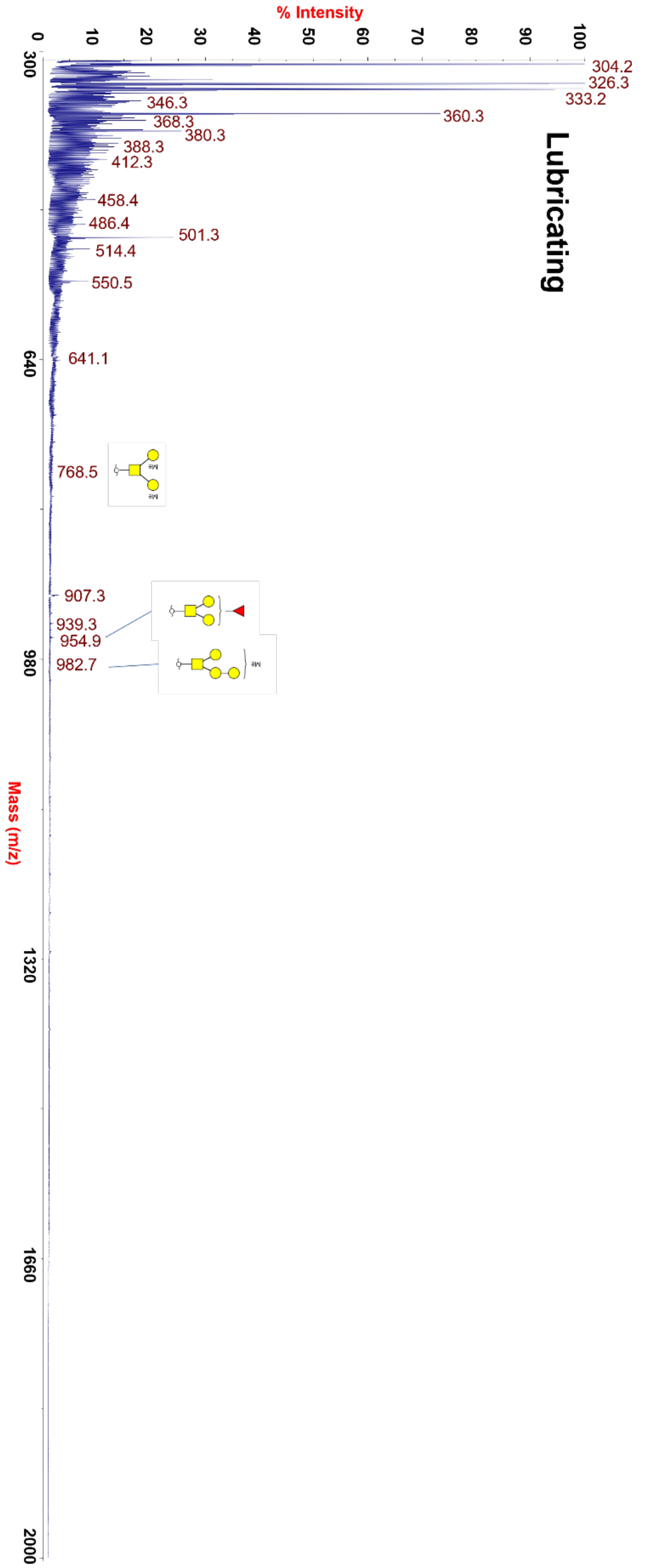

**Figure S10.** Glycomic mass spectrum showing m/z peaks and compositional assignments of extracted *O*-glycans from *C. aspersum* lubricating mucus samples. Samples were per-deuter-*O*-methylated and adducts shown are Na+.

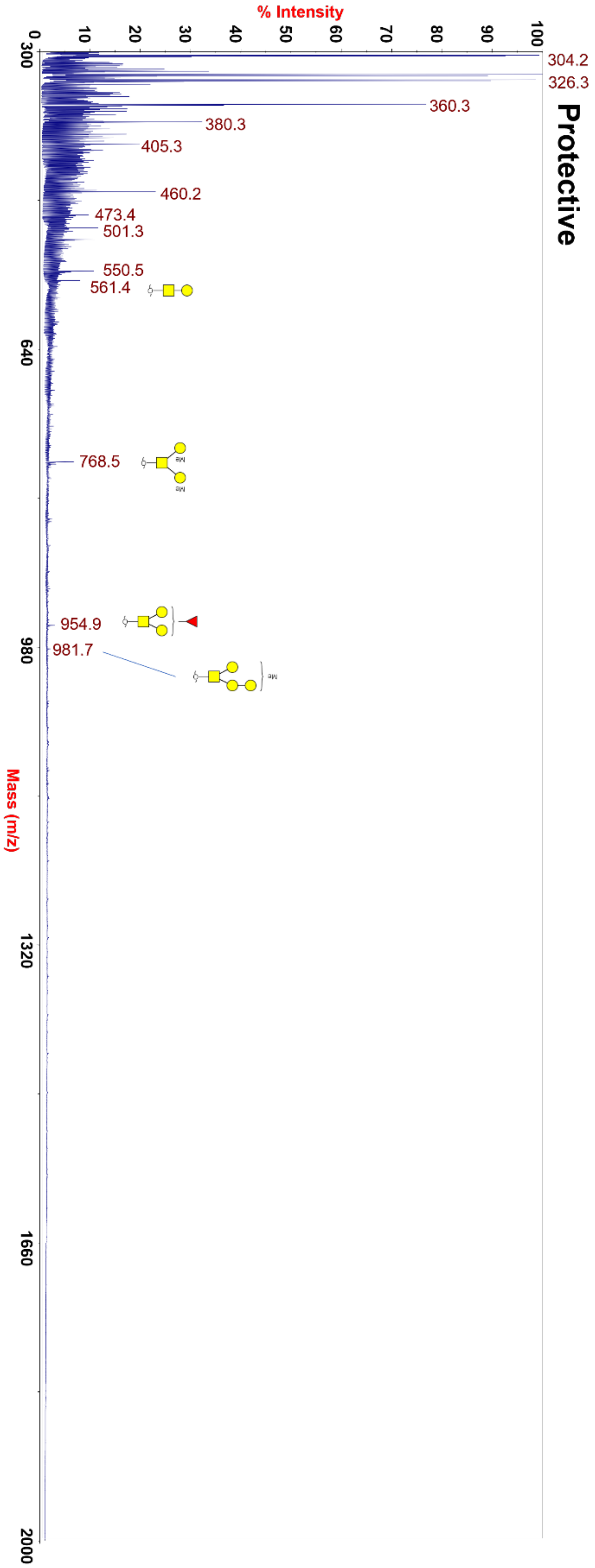

**Figure S11.** Glycomic mass spectrum showing m/z peaks and compositional assignments of extracted *O*-glycans from *C. aspersum* protective mucus samples. Insets show magnified region where low-intensity glycan peaks were found. Samples were per-deuter-*O*-methylated and adducts shown are Na+.

| Sample | Glycan m/z | Monosaccharide Composition | Proposed Structure | Relative Percentage |
|---|---|---|---|---|
| Adhesive | 561.5 | Gal1GalNAc1 | 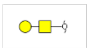 | 48.22% |
| | 768.7 | Gal2GalNAc1 + 2Me | 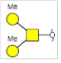 | 14.56% |
| | 771.7 | Gal2GalNAc1 + 1Me | 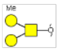 | 14.66% |
| | 774.7 | Gal2GalNAc1 | 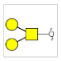 | 9.56% |
| | 981.9 | Gal3GalNAc1 + 1Me | 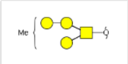 | 5.30% |
| | 1192.1 | NeuAc1Gal1HexNAc1GalNAc1 | 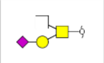 | 5.45% |
| | 1552.4 | NeuAc1Fuc2Gal1HexNAc1GalNAc1 | 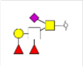 | 2.25% |
| Lubricating | 768.7 | Gal2GalNAc1 + 2Me | 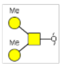 | 14.56% |
| | 954.97 | Fuc1Gal2GalNAc1 | 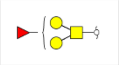 | 5.30% |
| | 982.8 | Gal3GalNAc1 + 1Me | 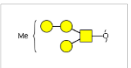 | 5.30% |
| Protective | 534.5 | Gal1GalNAc1 |  | 45.10% |
| | 738.8 | Gal2GalNAc1 + 2Me | 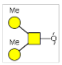 | 48.90% |
| | 942.8 | Fuc1Gal2GalNAc1 | 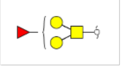 | 2.80% |
| | 983.9 | Gal3GalNAc1 + 1Me | 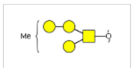 | 2.20% |

**Table S6.** *O*-Glycans extracted from adhesive, lubricating, and protective, *C. aspersum* snail mucus proteins that were detected via glycomic mass spectrometry analysis. Relative percentage refers to the ratio of the area of each individual glycan MS peak to the total area of all glycan peaks for a given experiment. Samples were per-deuter-*O*-methylated and adducts shown are Na+.

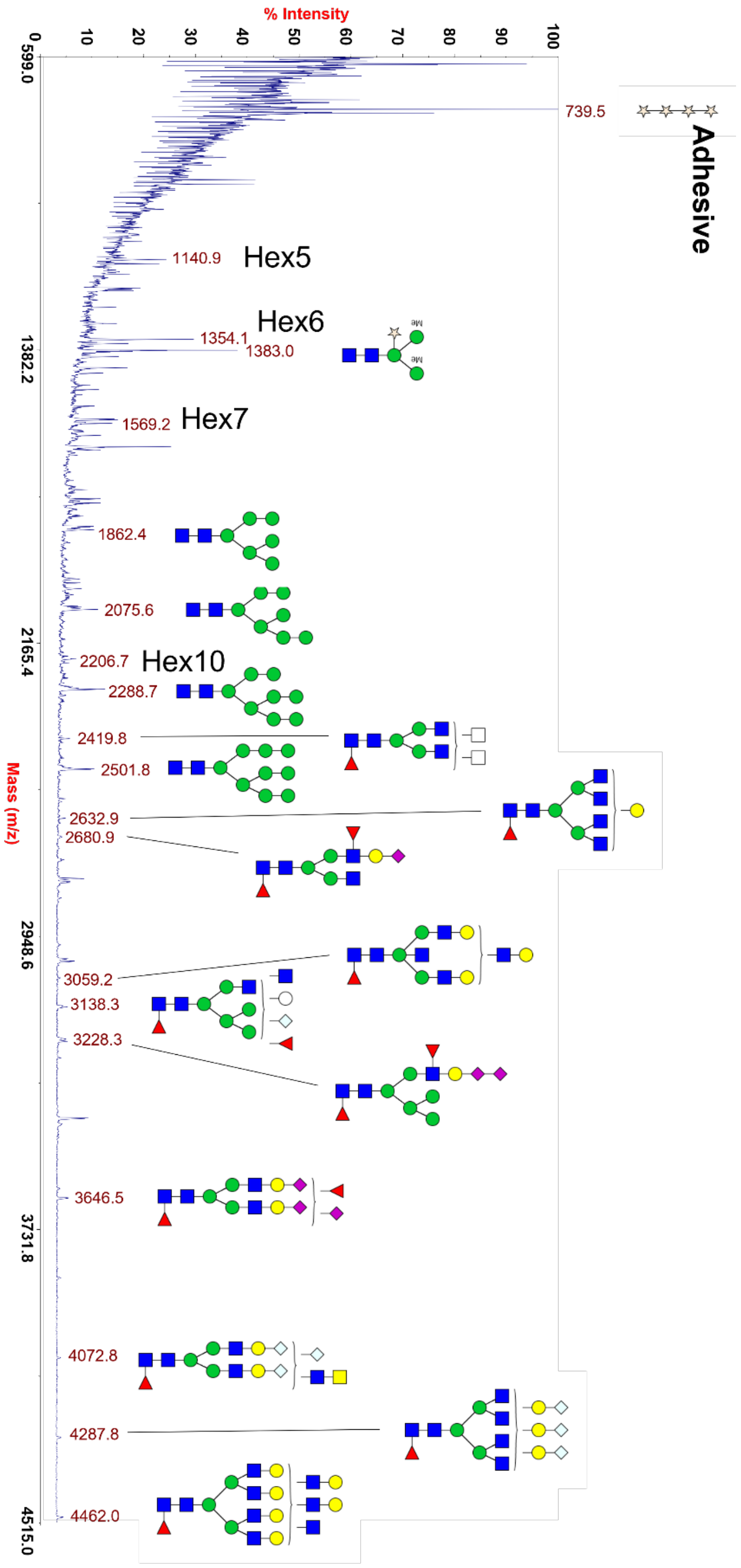


**Figure S12.** Glycomic mass spectrum showing m/z peaks and compositional assignments of extracted *N*-glycans from *C. aspersum* adhesive mucus samples. Samples were per-deuter-*O*-methylated and adducts shown are Na+.

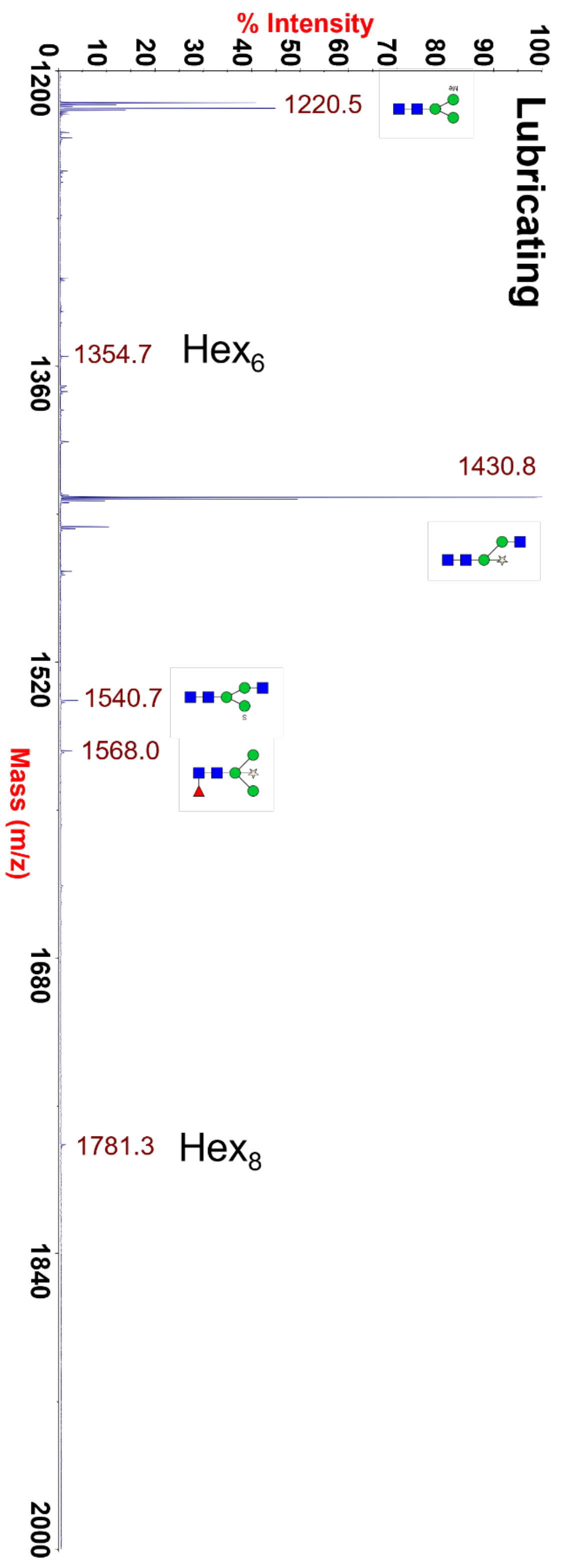


**Figure S13.** Glycomic mass spectrum showing m/z peaks and compositional assignments of extracted *N*-glycans from *C. aspersum* lubricating mucus samples. Samples were per-deuter-*O*-methylated and adducts shown are Na+.

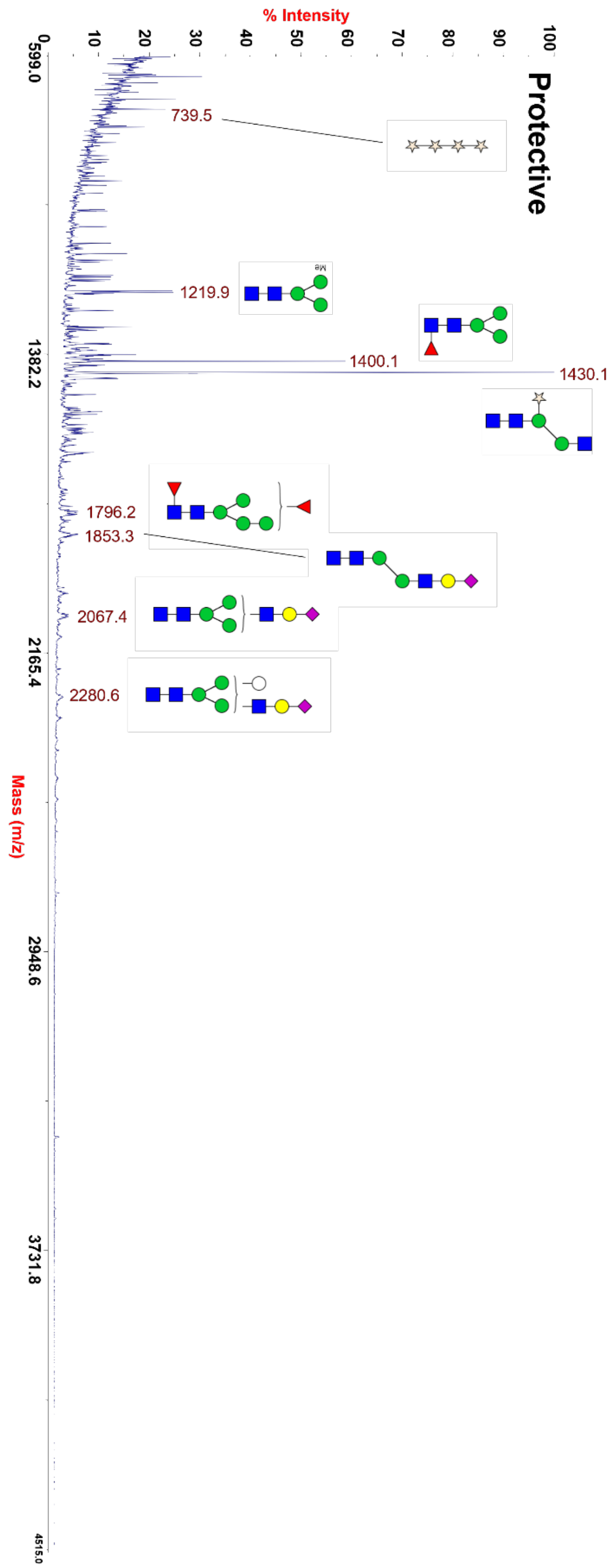


**Figure S14.** Glycomic mass spectrum showing m/z peaks and compositional assignments of extracted *N*-glycans from *C. aspersum* protective mucus samples. Samples were per-deuter-*O*-methylated and adducts shown are Na+.

| Sample | Glycan m/z | Monosaccharide Composition | Proposed Structure | Relative Percentage |
|---|---|---|---|---|
| Adhesive | 1382.96 | Man3GlcNAc2Xyl1 + 2 Me | 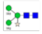 | 30% |
| | 1862.33 | Man6GlcNAc2 | 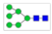 | 8% |
| | 2075.46 | Man7GlcNAc2 | 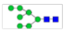 | 9% |
| | 2288.6 | Man8GlcNAc2 | 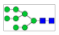 | 10% |
| | 2419.69 | HexNAc2GlcNAc2Man3GlcNAc2Fuc1 | 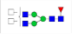 | 4% |
| | 2501.73 | Man9GlcNAc2 | 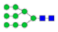 | 8% |
| | 2632.81 | Gal1GlcNAc4Man3GlcNAc2Fuc1 | 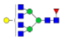 | 4% |
| | 2680.8 | NeuAc1Gal1GlcNAc2Man3GlcNAc2Fuc2 | 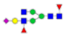 | 3% |
| | 3059.06 | Gal3GlcNAc4Man3GlcNAc2Fuc1 | 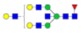 | 3% |
| | 3138.14 | Fuc1Hex1GlcNAc2Man5GlcNAc2Fuc1 | 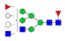 | 4% |
| | 3228.19 | NeuAc2Gal1GlcNAc1Man5GlcNAc2Fuc1 | 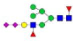 | 4% |
| | 3646.33 | NeuAc3Fuc1Gal2GlcNAc2Man3GlcNAc2Fuc1 | 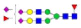 | 4% |
| | 4072.63 | GalNAc1NeuGc3Gal2GlcNAc3Man3GlcNAc2Fuc1 | 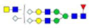 | 3% |
| | 4287.6 | NeuGc3Gal3GlcNAc4Man3GlcNAc2Fuc1 | 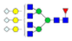 | 3% |
| | 4461.8 | Gal6GlcNAc7Man3GlcNAc2Fuc1 | 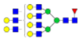 | 3% |
| Lubricating | 1219.92 | Man$_3$GlcNAc$_2$ + 1 Me | 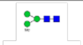 | 32% |
| | 1430.07 | GlcNAc$_1$Man$_2$GlcNAc$_2$Xyl$_1$ | 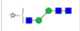 | 64% |
| | 1568 | Man$_3$GlcNAc$_2$Fuc$_1$Xyl$_1$ | 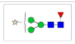 | 4% |
| Protective | 1219.92 | Man$_3$GlcNAc$_2$ + 1 Me | 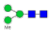 | 12% |
| | 1400.13 | Man$_3$GlcNAc$_2$Fuc$_1$ | 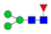 | 29% |
| | 1430.07 | GlcNAc$_1$Man$_2$GlcNAc$_2$Xyl$_1$ | 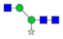 | 50% |
| | 1796.24 | Man$_4$GlcNAc$_2$Fuc$_2$ | 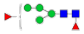 | 3% |
| | 1853.33 | NeuAc$_1$Gal$_1$GlcNAc$_1$Man$_2$GlcNAc$_2$ | 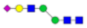 | 3% |
| | 2067.43 | NeuAc$_1$Gal$_1$GlcNAc$_1$Man$_3$GlcNAc$_2$ | 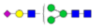 | 2% |
| | 2280.56 | NeuAc$_1$Gal$_1$Hex$_1$GlcNAc$_1$Man$_3$GlcNAc$_2$ | 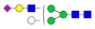 | 1% |

**Table S7.** *N*-Glycans extracted from adhesive, lubricating, and protective, *C. aspersum* snail mucus proteins that were detected via glycomic mass spectrometry analysis. Relative percentage refers to the ratio of the area of each individual glycan MS peak to the total area of all glycan peaks for a given experiment. Samples were per-deuter-*O*-methylated and adducts shown are Na+.

**5. Scanning electron microscopy**

**General methods.** The samples were made by letting the snail to crawl and leave mucus on an aluminum SEM pin stubs, similarly to the silicon wafer samples for AFM, and air-dried overnight. The samples were sputter-coated with gold to a thickness of 5 nm using a Leica EM ACE600 Coater for better electrical conductivity. These samples were then imaged in a Thermo Scientific (FEI) Helios NanoLab 660 FIB-SEM with HT of 5 kV, current of 6.3, 13 and 25 pA with ETD (Everhart-Thornley) detector. EDS (energy-dispersive X-ray spectroscopy) mapping was collected with an Oxford detector at HT of 10 kV and current of 1.6 nA. Data was collected and analyzed using AZtec software.[25]

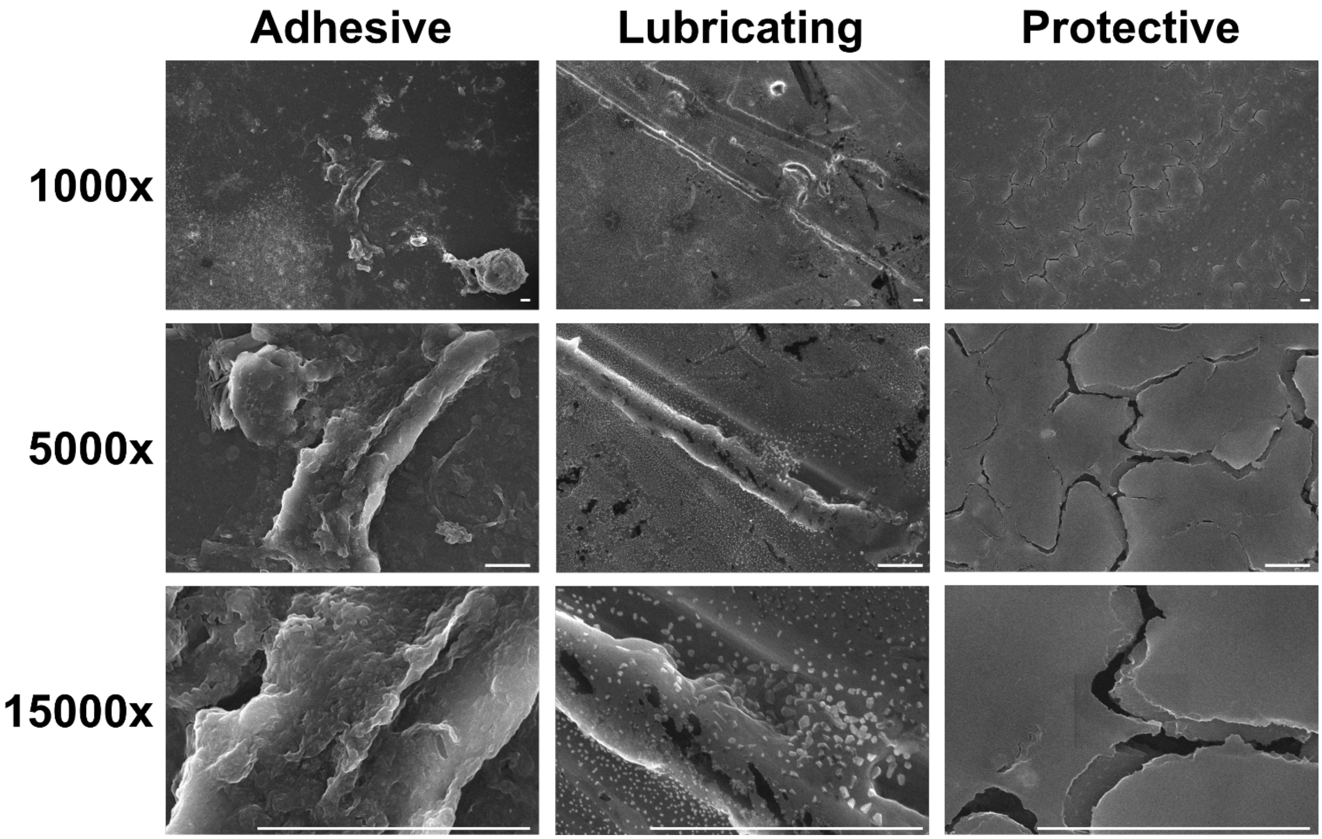


**Figure S15.** SEM imaging of *C. aspersum* adhesive, lubricating, and protective mucus at several magnification levels. Scale bars represent 6 μm.

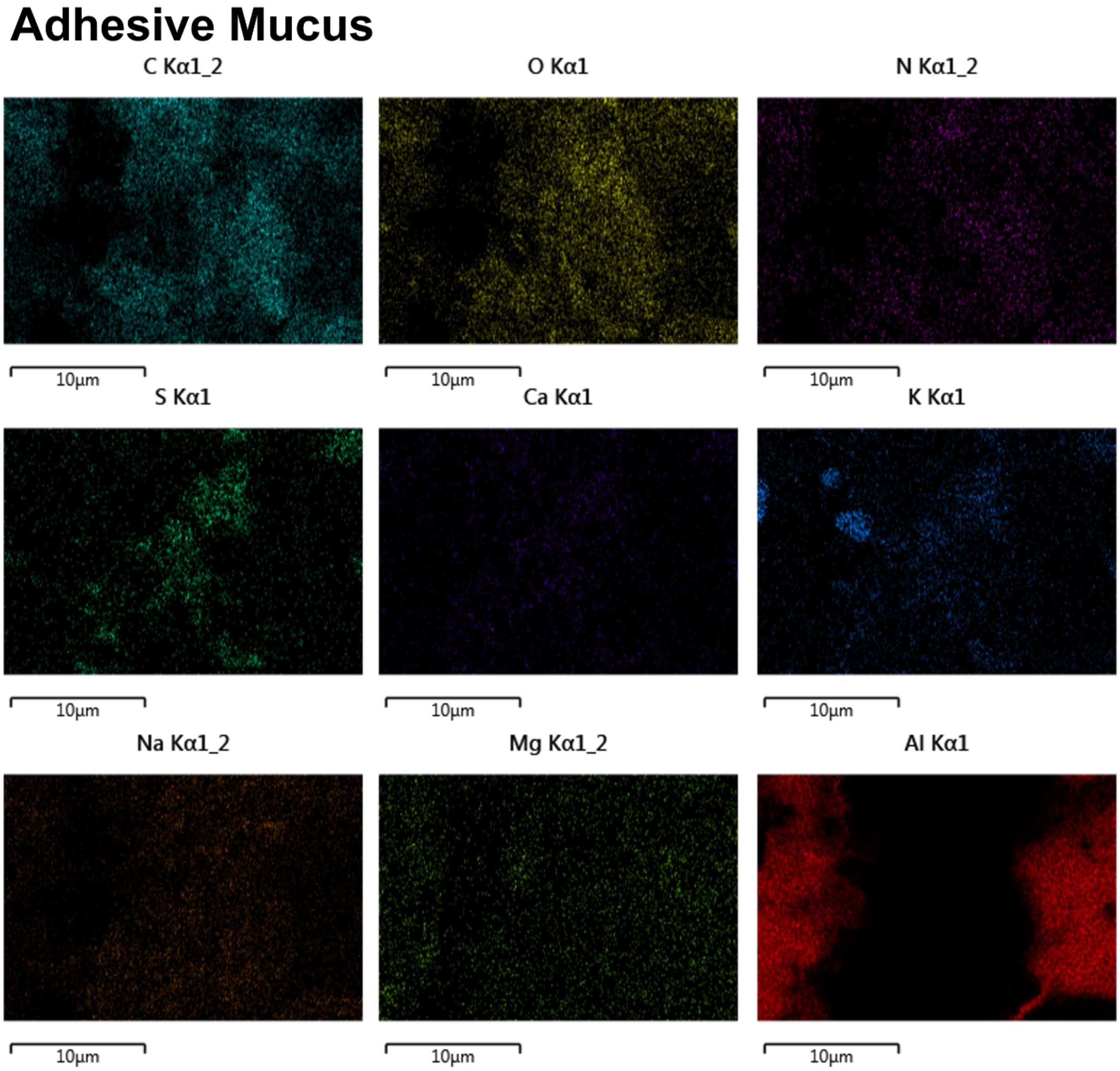


**Figure S16.** SEM EDX overlay images of *C. aspersum* adhesive snail mucus showing localization of detected elements. Scale bars shown.

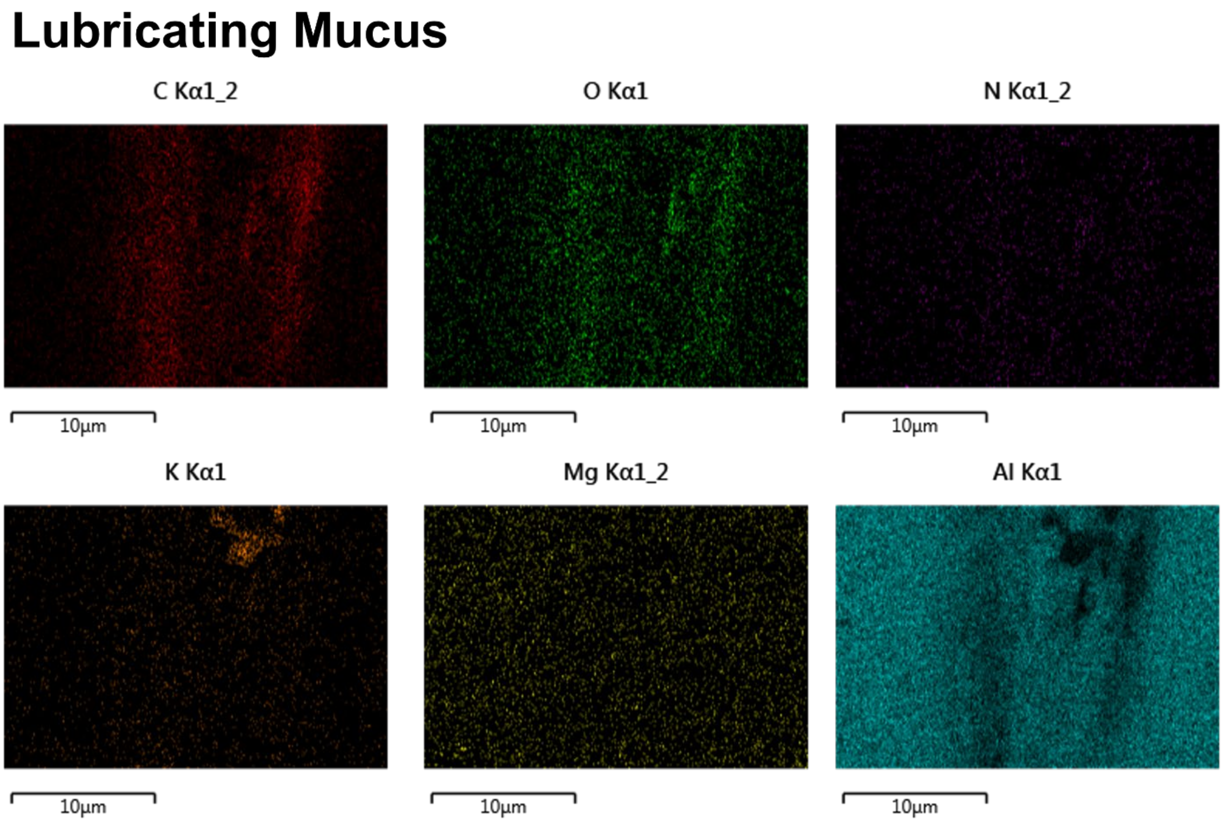


**Figure S17.** SEM EDX overlay images of *C. aspersum* lubricating snail mucus showing localization of detected elements. Scale bars shown.

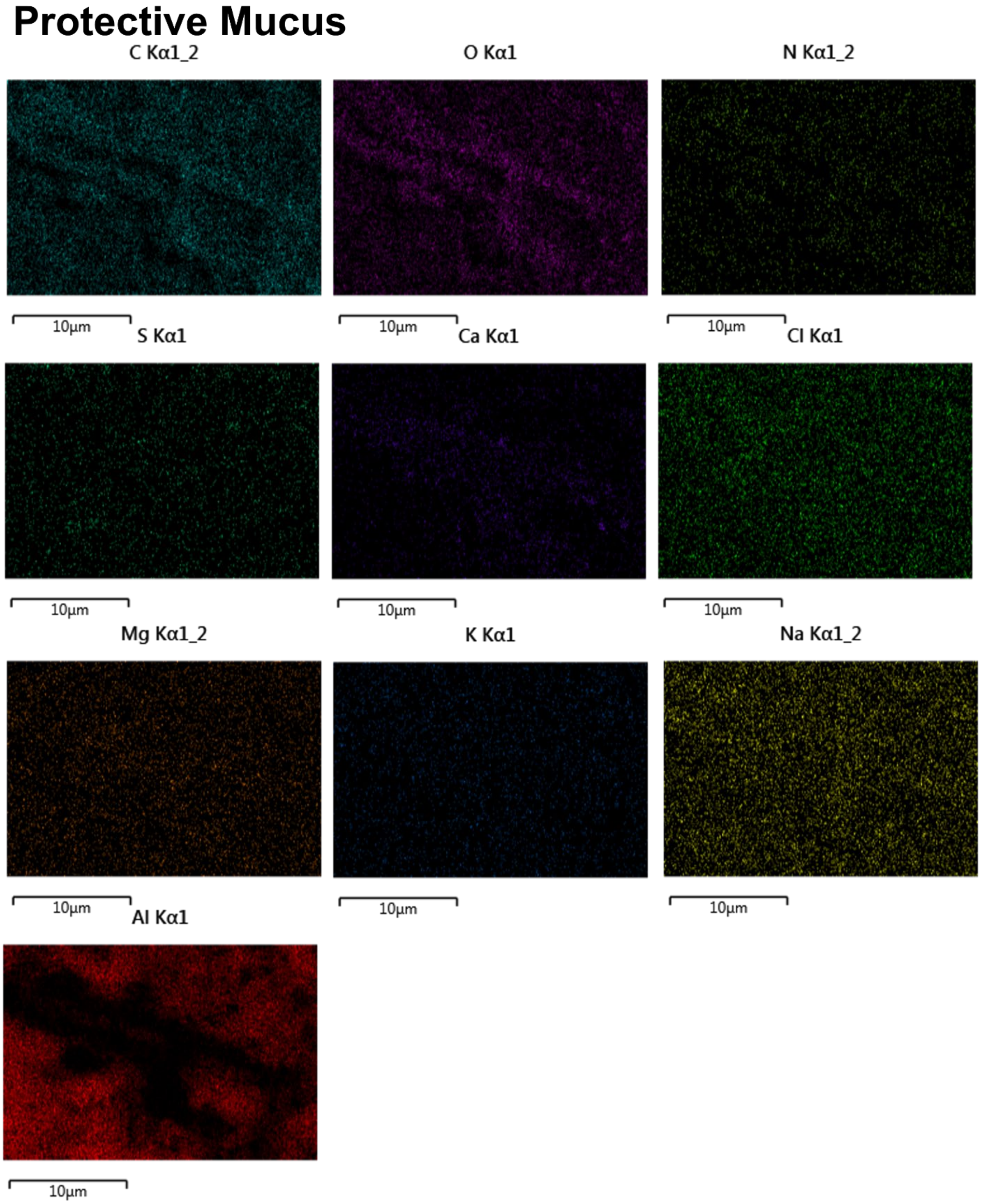


**Figure S18.** SEM EDX overlay images of *C. aspersum* protective snail mucus showing localization of detected elements. Scale bars shown.

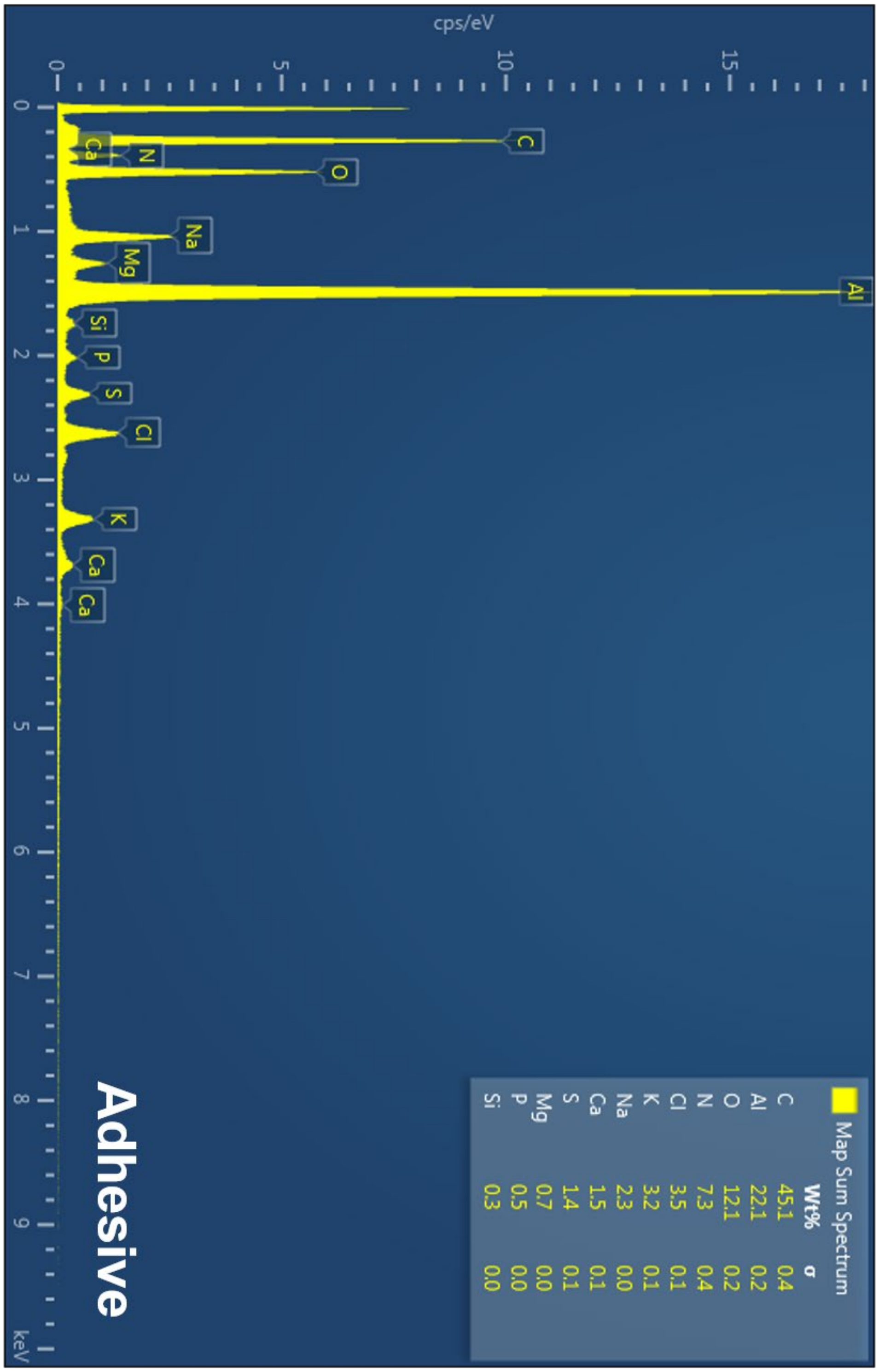


**Figure S19.** Energy Dispersive X-Ray (EDX) spectra showing elemental composition of *C. aspersum* adhesive mucus.

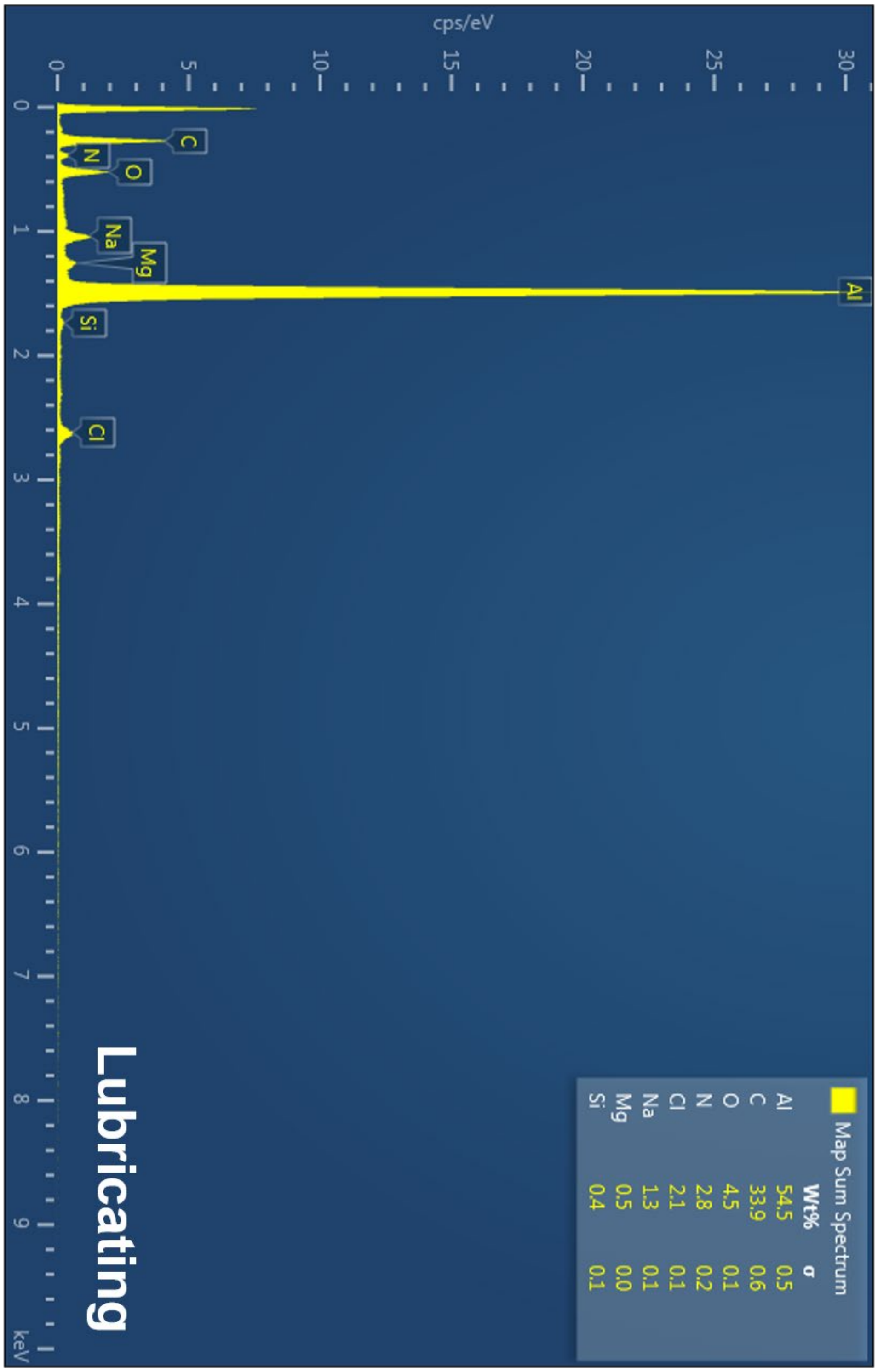


**Figure S20.** Energy Dispersive X-Ray (EDX) spectra showing elemental composition of *C. aspersum* lubricating mucus.

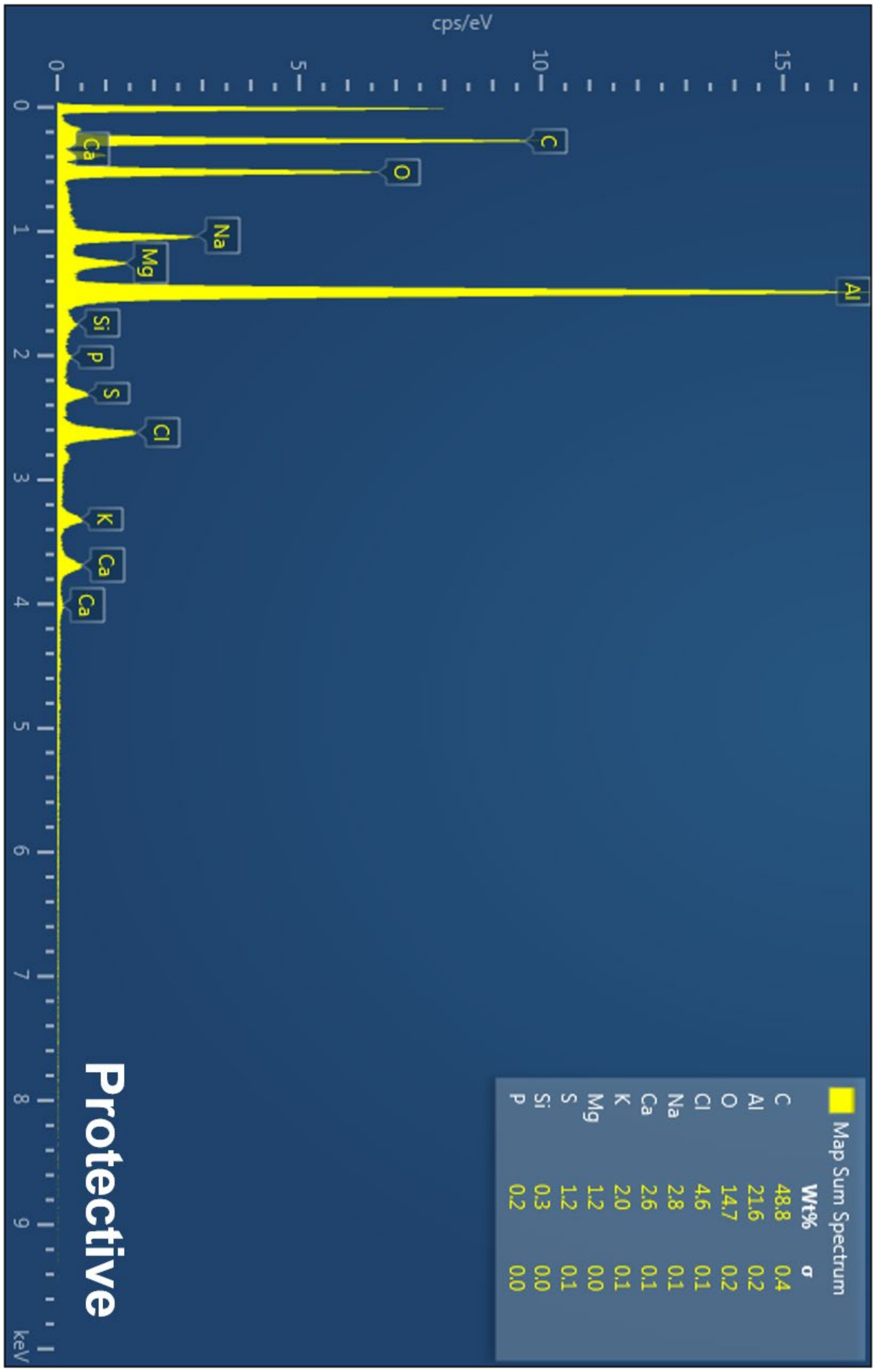


**Figure S21.** Energy Dispersive X-Ray (EDX) spectra showing elemental composition of *C. aspersum* protective mucus.

### 6. Atomic force microscopy

#### General methods.

*AFM topography*

The sample was made by letting the snail to crawl and leave mucus on a silicon wafer, which was subsequently scanned by using an AFM (Multimode 8, Bruker). Mucus topographies were imaged using the AFM tapping mode with a probe (SCANASYST-AIR, Bruker) that has a tip radius of 20 nm. To locate positions of interest, we used a microscope to find sample features that are clean and intact, made the AFM probe to be above the feature, and then lowered the AFM probe to enable tip-sample interaction that is required for topography measurement. We controlled the scan size to be 10-20 um depending on the feature sizes, scan rate to be 0.8 Hz, and the pixel number to be 256 x 256. Each sample was scanned for more than 5 topographies images to avoid site selection bias and to ensure sample characteristics are captured.

*Nano-indentation experimental procedure*

The stiffness of mucus samples was characterized using the AFM nano-indentation method,[26] where an indenter (MLCT-E, Bruker) with radius of 20 nm and a spring constant of 0.139 N/m was used. The indentation deflection sensitivity was 40.7 nm/V, calibrated by performing an indentation on the silicon wafer substrate. To select the points for indentations, we first used the AFM tapping mode to map the surface materials, which were subsequently grouped into 3 to 5 categories based on their size and shape. For example, aggregates having sharp edges and have the size of ~1 um are regarded as one type; the material that spans across the entire material map and supports all kinds of aggregates is regarded as the other type. After we identified the material types on the sample, we offset the scanning tip to the peak of certain material type and decreased the scan size to 10 nm x 10 nm before we switched the AFM mode from the tapping mode to the indentation mode. The indentation rate was set at 1 Hz and the tip deflection signal, which triggers the approaching movement to switch to the retracting movement, was tuned to increase from 0.05 V until the voltage that leads to an indentation depth of ~2 nm, of which the indentation profiles were collected three times for analysis (Methods). Three locations of each material type were characterized to obtain statistically reliable stiffness results.

*Stiffness and work of adhesion characterization via the JKR model*

The stiffness of mucus samples was characterized using AFM nano-indentation method,[26] where an indenter (MLCT-E, Bruker) with radius of 20 nm and a spring constant of 0.139 N/m was used. The indentation deflection sensitivity was 40.7 nm/V, calibrated by performing an indentation on the silicon wafer substrate. Peaks of three mucus aggregates are indented to obtain the force *vs.* displacement relationships, of which the retracting portion of the indenting profiles were subsequently analyzed by using the Johnson–Kendall–Roberts (JKR) model, given by

$$E_{\mathrm{JKR}} = \frac{9\pi \mathrm{R}^2 \Delta r}{2a_0^3},$$

$$P_{\mathrm{adh}} = -\frac{3}{2}\pi \Delta r \mathrm{R},$$

$$h_t - h_0 = \frac{a_0^2}{\mathrm{R}}\left(\frac{1+\sqrt{1-\frac{P}{P_{\mathrm{adh}}}}}{2}\right)^{\frac{4}{3}} - \frac{2}{3}\frac{a_0^2}{\mathrm{R}}\left(\frac{1+\sqrt{1-\frac{P}{P_{\mathrm{adh}}}}}{2}\right)^{\frac{1}{3}},$$

where $E_{JKR}$ is the Young's modulus, R is the tip radius, $\Delta r$ is the work of adhesion, $a_0$ is the contact area when the contract force is zero, $P_{adh}$ is the pull-off force, $h_t$ is the indentation depth, $h_0$ is the contact point where the pull-off force shows, and $P$ is the load. The work of adhesion was measured by the area enclosed by the approaching and the retracting indentation force-displacement curves, and was normalized by the probe sample contact area ($a_0$), given by

$a_0 = \pi R h_t$.

Force-retract curves were selected for analysis if $R^2$ values were greater than .96 and $E$ and $W$ had non-negative values.

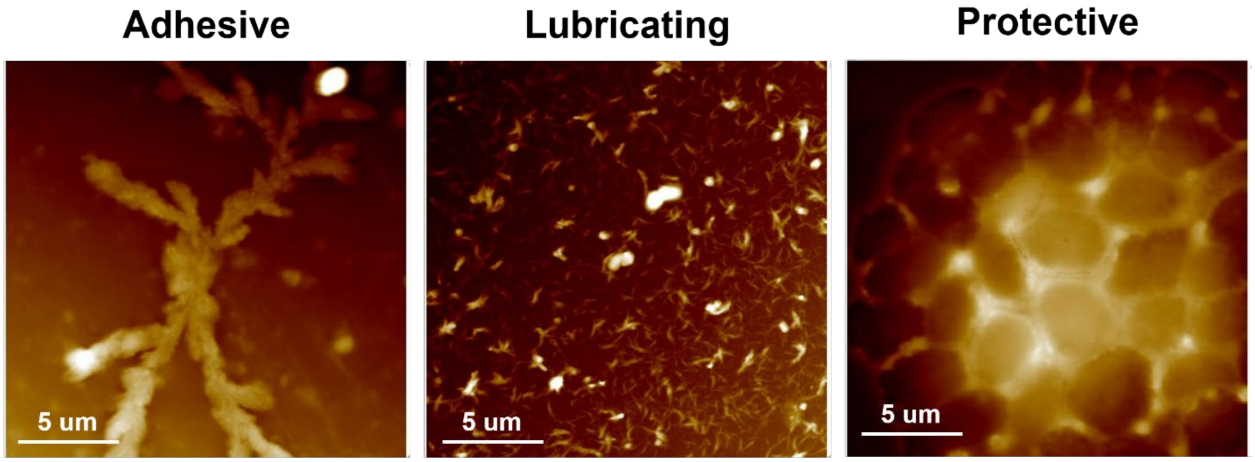

**Figure S22.** AFM topography imaging. Scale bars shown.

### References


1 Newar, J. & Ghatak, A. Studies on the adhesive property of snail adhesive mucus. *Langmuir* **31**, 12155-12160 (2015).

2 Greistorfer, S. *et al.* Snail mucus− glandular origin and composition in Helix pomatia. *Zoology* **122**, 126-138 (2017).

3 Ballance, S. *et al.* Partial characterisation of high-molecular weight glycoconjugates in the trail mucus of the freshwater pond snail Lymnaea stagnalis. *Comparative Biochemistry and Physiology Part B: Biochemistry and Molecular Biology* **137**, 475-486 (2004).

4 Corfield, A. P. *Glycoprotein methods and protocols: The mucins*. Vol. 125 (Springer Science & Business Media, 2000).

5 Kilcoyne, M., Gerlach, J. Q., Farrell, M. P., Bhavanandan, V. P. & Joshi, L. Periodic acid–Schiff's reagent assay for carbohydrates in a microtiter plate format. *Analytical biochemistry* **416**, 18-26 (2011).

6 Dheilly, N. M. *et al.* A family of variable immunoglobulin and lectin domain containing molecules in the snail Biomphalaria glabrata. *Developmental & Comparative Immunology* **48**, 234-243, doi:https://doi.org/10.1016/j.dci.2014.10.009 (2015).

7 Barcia, R., Lopez-García, J. M. & Ramos-Martínez, J. I. The 28S fraction of rRNA in molluscs displays electrophoretic behaviour different from that of mammal cells. *IUBMB Life* **42**, 1089-1092 (1997).

8 Brown, J., Pirrung, M. & McCue, L. A. FQC Dashboard: integrates FastQC results into a web-based, interactive, and extensible FASTQ quality control tool. *Bioinformatics* **33**, 3137-3139 (2017).

9 Bolger, A. M., Lohse, M. & Usadel, B. Trimmomatic: a flexible trimmer for Illumina sequence data. *Bioinformatics* **30**, 2114-2120 (2014).

10 Haas, B. J. *et al.* De novo transcript sequence reconstruction from RNA-seq using the Trinity platform for reference generation and analysis. *Nature protocols* **8**, 1494-1512 (2013).

11 Davidson, N. M., Hawkins, A. D. & Oshlack, A. SuperTranscripts: a data driven reference for analysis and visualisation of transcriptomes. *Genome biology* **18**, 1-10 (2017).

12 Helsens, K., Martens, L., Vandekerckhove, J. & Gevaert, K. MascotDatfile: an open-source library to fully parse and analyse MASCOT MS/MS search results. *Proteomics* **7**, 364-366 (2007).

13 Searle, B. C. Scaffold: a bioinformatic tool for validating MS/MS-based proteomic studies. *Proteomics* **10**, 1265-1269 (2010).

14 Liu, W. *et al.* Stress-Induced Mucus Secretion and Its Composition by a Combination of Proteomics and Metabolomics of the Jellyfish Aurelia coerulea. *Marine Drugs* **16**, 341 (2018).

15 Espinosa, E. P., Koller, A. & Allam, B. Proteomic characterization of mucosal secretions in the eastern oyster, Crassostrea virginica. *Journal of proteomics* **132**, 63-76 (2016).

16 Ballard, K. R., Klein, A. H., Hayes, R. A., Wang, T. & Cummins, S. F. The protein and volatile components of trail mucus in the Common Garden Snail, Cornu aspersum. *PloS one* **16**, e0251565 (2021).

17 Johnson, M. *et al.* NCBI BLAST: a better web interface. *Nucleic acids research* **36**, W5-W9 (2008).

18 Thompson, J. D., Gibson, T. J. & Higgins, D. G. Multiple sequence alignment using ClustalW and ClustalX. *Current protocols in bioinformatics*, 2.3. 1-2.3. 22 (2003).

19 Letunic, I. & Bork, P. Interactive Tree Of Life (iTOL) v5: an online tool for phylogenetic tree display and annotation. *Nucleic acids research* **49**, W293-W296 (2021).

20 Potter, S. C. *et al.* HMMER web server: 2018 update. *Nucleic acids research* **46**, W200-W204 (2018).

21 Procter, J. B. *et al.* in *Multiple Sequence Alignment* 203-224 (Springer, 2021).

22 Kang, P., Mechref, Y., Kyselova, Z., Goetz, J. A. & Novotny, M. V. Comparative glycomic mapping through quantitative permethylation and stable-isotope labeling. *Analytical chemistry* **79**, 6064-6073 (2007).
23 Wu, Y. *et al.* N-Glycomic profiling reveals dysregulated glycans related to oral cancer using MALDI-MS. *Analytical and Bioanalytical Chemistry* **414**, 1881-1890 (2022).
24 Damerell, D. *et al.* in *Glycoinformatics* 3-15 (Springer, 2015).
25 Burgess, S. & Pinard, P. AZtec Wave–a New Way to Achieve Combined EDS and WDS Capability on SEM. *Microscopy and Microanalysis* **26**, 114-115 (2020).
26 Wu, G., Gotthardt, M. & Gollasch, M. Assessment of nanoindentation in stiffness measurement of soft biomaterials: kidney, liver, spleen and uterus. *Scientific reports* **10**, 1-11 (2020).
